# Heterochromatin fidelity is a therapeutic vulnerability in lymphoma and other human cancers

**DOI:** 10.1101/2025.01.31.635709

**Authors:** Mohamad Ali Najia, Deepak K. Jha, Cheng Zhang, Benoit Laurent, Caroline Kubaczka, Arianna Markel, Christopher Li, Vivian Morris, Allison Tompkins, Luca Hensch, Yue Qin, Bjoern Chapuy, Yu-Chung Huang, Michael Morse, Matthew R. Marunde, Anup Vaidya, Zachary B. Gillespie, Sarah A. Howard, Trista E. North, Daniel Dominguez, Michael-Christopher Keogh, Thorsten M. Schlaeger, Yang Shi, Hu Li, Margaret M. Shipp, Paul C. Blainey, George Q. Daley

## Abstract

Genes involved in the regulation of chromatin structure are frequently disrupted in cancer, contributing to an aberrant transcriptome and phenotypic plasticity. Yet, therapeutics targeting mutant forms of chromatin-modifying enzymes have yielded only modest clinical utility, underscoring the difficulty of targeting the epigenomic underpinnings of aberrant gene regulatory networks. Here, we sought to identify novel epigenetic vulnerabilities in diffuse large B-cell lymphoma (DLBCL). Through phenotypic screens and biochemical analysis, we demonstrated that inhibition of the H3K9 demethylases KDM4A and KDM4C elicits potent, subtype-agnostic cytotoxicity by antagonizing transcriptional networks associated with B-cell identity and epigenetically rewiring heterochromatin. KDM4 demethylases associated with the KRAB zinc finger ZNF587, and their enzymatic inhibition led to DNA replication stress and DNA damage-induced cGAS-STING activation. Broad surveys of transcriptional data from patients also revealed KDM4 family dysregulation in several other cancer types. To explore this potential therapeutic avenue, we performed high-throughput small molecule screens with H3K9me3 nucleosome substrates and identified novel KDM4 demethylase inhibitors. AI-guided protein-ligand binding predictions suggested diverse modes of action for various small molecule hits. Our findings underscore the relevance of targeting fundamental transcriptional and epigenetic mechanisms for anti-cancer therapy.

**HIGHLIGHTS:**

- Phenotypic screens identified JIB-04 as a potent anti-cancer agent for multiple subtypes of diffuse large B-cell lymphoma
- JIB-04 binds and inhibits KDM4 demethylases resulting in epigenomic rewiring of heterochromatin
- KDM4 demethylases cooperate with KRAB zinc fingers to limit DNA replication stress, and KDM4 inhibition instigates DNA-damage and cGAS-STING activation in several human cancers
- High-throughput small molecule screens with semi-synthetic nucleosome substrates and AI-guided molecular docking simulations identify novel KDM4 inhibitors

## INTRODUCTION

Cancer cells must continuously respond and adapt to environmental and selective pressures. Chromatin is the medium by which multiple, upstream signaling inputs converge to drive or repress gene expression programs^1–3^. Chromatin modulators elicit, maintain and tune transcriptional control through a diverse compendium of covalent modifications on histones and DNA^4,5^. These modifications are capable of altering chromatin structure to usher transcriptionally active euchromatin or inhibitory heterochromatin^6–9^. The regulation of post-translational modifications on histones, and chromatin structure more generally, are thus integral for fundamental cellular processes, such as transcription^10,11^, DNA repair^12^, genome stability^13^ and cell cycle regulation^14^.

Unsurprisingly, genes involved in chromatin and transcriptional modulation are frequently mutated, deleted or amplified in various types of human cancers^15–19^. The oncogenic cooption of chromatin machinery can consequently disrupt chromatin states and result in aberrant gene regulation^20–23^. Clinically, these molecular mechanisms manifest to influence the phenotypic plasticity, metastatic capacity, proliferative nature and therapeutic susceptibility of tumors^15,24–30^. However, therapeutic strategies to target the epigenomic underpinnings of aberrant gene regulatory networks in human malignancies has proven a major challenge.

Diffuse large B-cell lymphoma (DLBCL) can be stratified on transcriptional states with prognostic significance^31–35^, and is an ideal model to study chromatin mechanisms of oncogenic states^36,37^. The germinal center B-cell (GCB) subtype exhibits transcriptional signatures reflective of the germinal center reaction and is characterized by higher overall patient survival compared to activated B cell (ABC) subtypes, which transcriptionally resemble plasmacytic lineages^31,33,34,38^. Of note, lineage-specific transcription factors (TFs: e.g., *IRF4, ETV6, EBF1, IRF8*), signaling pathways and effectors (e.g. *CD70, CD79B, MYD88, JAK3, PI3KCD*), and epigenetic factors (e.g. *EZH2, CREBBP, EP300, KMT2C, KMT2*D) are frequently mutated in patients^33,34,38–42^. These clinical observations have motivated proposals to target dysregulated chromatin pathways with mechanistic understanding and therapeutic intent. In this manner, EZH2 inhibitors are active against GCB subtypes, but not ABC subtypes^43–45^, potentially reflecting the importance of EZH2 during germinal center differentiation^46^. Despite significant efforts to pharmacologically target the EZH2 methyltransferase, EP300 and CREBBP acetyltransferases, and BRD4 acetyl-reader, clinical outcomes have been met with variable degrees of success^42–45,47,48^. The modest therapeutic benefits likely underscore an incomplete understanding of how various epigenetic factors impact cancer-specific genetic alterations and functionally interact with other oncogenic mechanisms. Furthermore, while combination cytotoxic chemotherapy and, more recently, adoptive T cell therapies have demonstrated remarkable promise in DLBCL, 30% of patients relapse or are refractory to first line therapies^49,50^, necessitating additional therapeutic strategies.

In this work, we sought to identify epigenetic, and thus potential therapeutic, vulnerabilities in DLBCL. Phenotypic screens with small molecule libraries targeting chromatin factors identified JIB-04, a pan-Jumonji domain inhibitor, as a highly potent anti-cancer agent for GCB and ABC subtypes of DLBCL. Mechanistically, JIB-04 binds the KDM4A and KDM4C histone H3K9 demethylases, resulting in genome-wide epigenomic rewiring, particularly in heterochromatic satellite regions, and cytotoxicity through DNA damage-induced cell cycle replication stress. Given the relevance of KDM4 proteins as anti-cancer therapeutic targets, we performed high-throughput small molecule enzymatic screens with H3K9me3 nucleosome substrates and identified several novel inhibitors. AI-guided, protein-ligand simulations suggested the binding sites of small molecule hits in KDM4 proteins and implicated diverse modes of enzymatic inhibition. Our work further advances therapeutic strategies for DLBCL and emphasizes the relevance of targeting fundamental transcriptional and epigenetic mechanisms for anti-cancer therapy.

## RESULTS

### Phenotypic screens identify novel anti-cancer compounds effective against DLBCL

To systematically search for epigenetic vulnerabilities in DLBCL, we performed phenotypic screens using a library of 145 small molecule compounds known to modulate the activity of a variety of chromatin “writer”, “reader” and “eraser” proteins (**Figure 1A**). In complementary screens, HBL-1 and OCI-Ly1 cells (representing ABC and GCB subtypes of DLBCL, respectively) were treated for five days with compounds at various doses (100 nM, 500 nM and 1 μM), and cell proliferation was assessed with CellTiter-Glo (CTG) assays. Compounds were defined as statistically significant hits if there was at least a two-fold reduction in CTG signal (FDR-adjusted *P* value < 0.0001) relative to DMSO vehicle controls (**Figure S1A-B, Supplemental Tables S1A-C**). At the 1 μM screening dose, 63% of hits (24 compounds) impacted both cell lines, suggesting conserved epigenetic vulnerabilities across DLBCL subtypes (**Figure 1B**). In contrast, eight compounds were hits only in HBL-1 cells and six compounds were hits only in OCI-Ly1 cells (**Figure 1B-C**). Reassuringly, Panobinostat, a histone deacetylase (HDAC) inhibitor and candidate to treat B cell malignancies^51^, was a hit in both cell lines, confirming the ability of our screens to identify agents with known anti-cancer activity. We identified 10 other HDAC inhibitors as hits in both cell lines, but due to the limited clinical efficacy of this drug class^52–57^, we opted to interrogate alternate hits from our screens.

**Figure 1.**
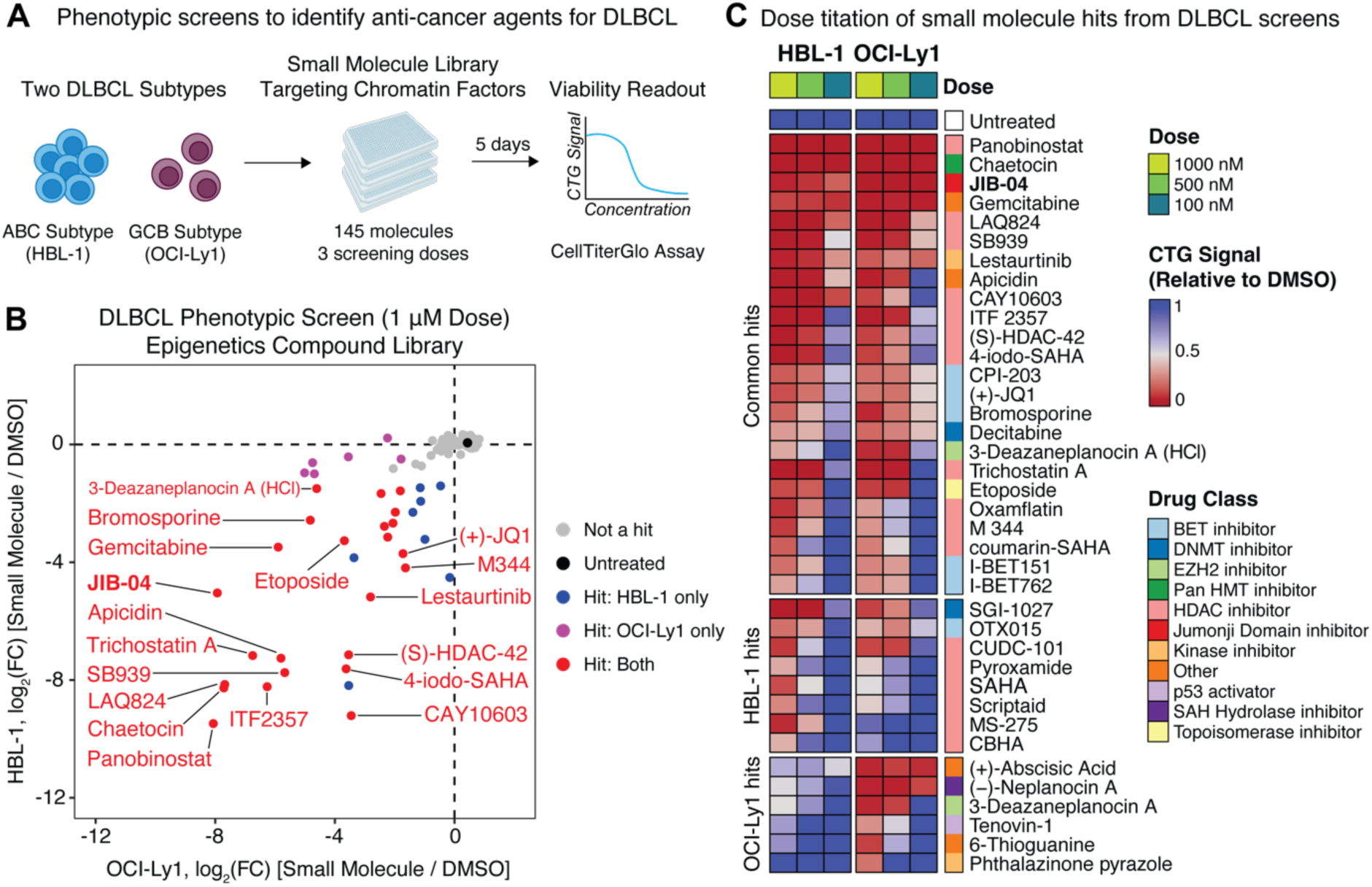
Phenotypic small molecule screens identify novel anti-cancer agents for DLBCL. **A)** Experimental scheme to identify compounds with anti-cancer activity against DLBCL via arrayed phenotypic screens. HBL-1 (ABC subtype) and OCI-Ly1 (GCB subtype) cells were treated with a library of 145 small molecules targeting chromatin factors at multiple doses (100 nM, 500 nM and 1 μM) for five days and cell proliferation was measured via CellTiter-Glo (CTG) luminescence assays. All compounds were screened with three replicates per dose. **B)** Fold change in CTG signal (1 μM of compounds compared to 0.01% DMSO vehicle controls) for HBL-1 and OCI-Ly1 cells after five days of treatment. Each datapoint represents the average fold-change in CTG signal per compound across all screen replicates. Compounds exhibiting a log_2_(fold change) < −1 and an FDR-adjusted *P* value < 0.0001 were defined as hits. **C)** Cytotoxicity of hits across all screening doses in HBL-1 and OCI-Ly1 cells. Compounds are rank ordered by CTG signal relative to DMSO on day 5 after treatment and grouped by conserved hits across both cell lines (top), hits only in HBL-1 cells (middle), and hits only in OCI-Ly1 cells (bottom). Relative CTG signal is the average of three screening replicates.

We further refined the prioritization of hits based on the potency across the doses screened. Of the 24 hits at the 1 μM dose, 5 compounds also scored as hits at 100 nM in both cell lines. Notably, JIB-04, a pan-Jumonji domain inhibitor, was one of the top hits across all doses and in both DLBCL subtypes (**Figure 1B-C**). The cytotoxicity exhibited by JIB-04 was more pronounced than inhibitors of EZH2 (GSK343, UNC1999 and 3-Deazaneplanocin A) and bromodomains (I-BET762, I-BET151, and (+)-JQ1) (**Figure S1C**). To the best of our knowledge, JIB-04 has not been studied within lymphoma or B-cell malignancies, which motivated us to focus on this potentially novel drug class for DLBCL. Collectively, our phenotypic screens confirmed known compounds with anti-cancer activity in lymphoma and identified JIB-04 as a novel agent with potent, subtype-agnostic cytotoxicity in DLBCL.

### JIB-04 exhibits potent activity across a broad compendium of DLBCL cell lines

We next validated JIB-04 as a hit from our primary screens in a further refined time and dose regimen, identifying a 99% reduction in CTG signal within two days of treatment at doses as low as 100 nM (**Figure 2A**). Importantly, JIB-04 exists in two stereoisomers^58^, and lymphoma cells were 100-fold more sensitive to the active (*E*) isomer than the inactive (*Z*) isomer (*P* < 0.01, **Figure S2A**). Furthermore, we verified that JIB-04 was eliciting cytotoxicity by induction of apoptosis through Annexin V positivity (**Figure S2B**) and cleaved PARP and caspase 3 (**Figure S2C**).

**Figure 2.**
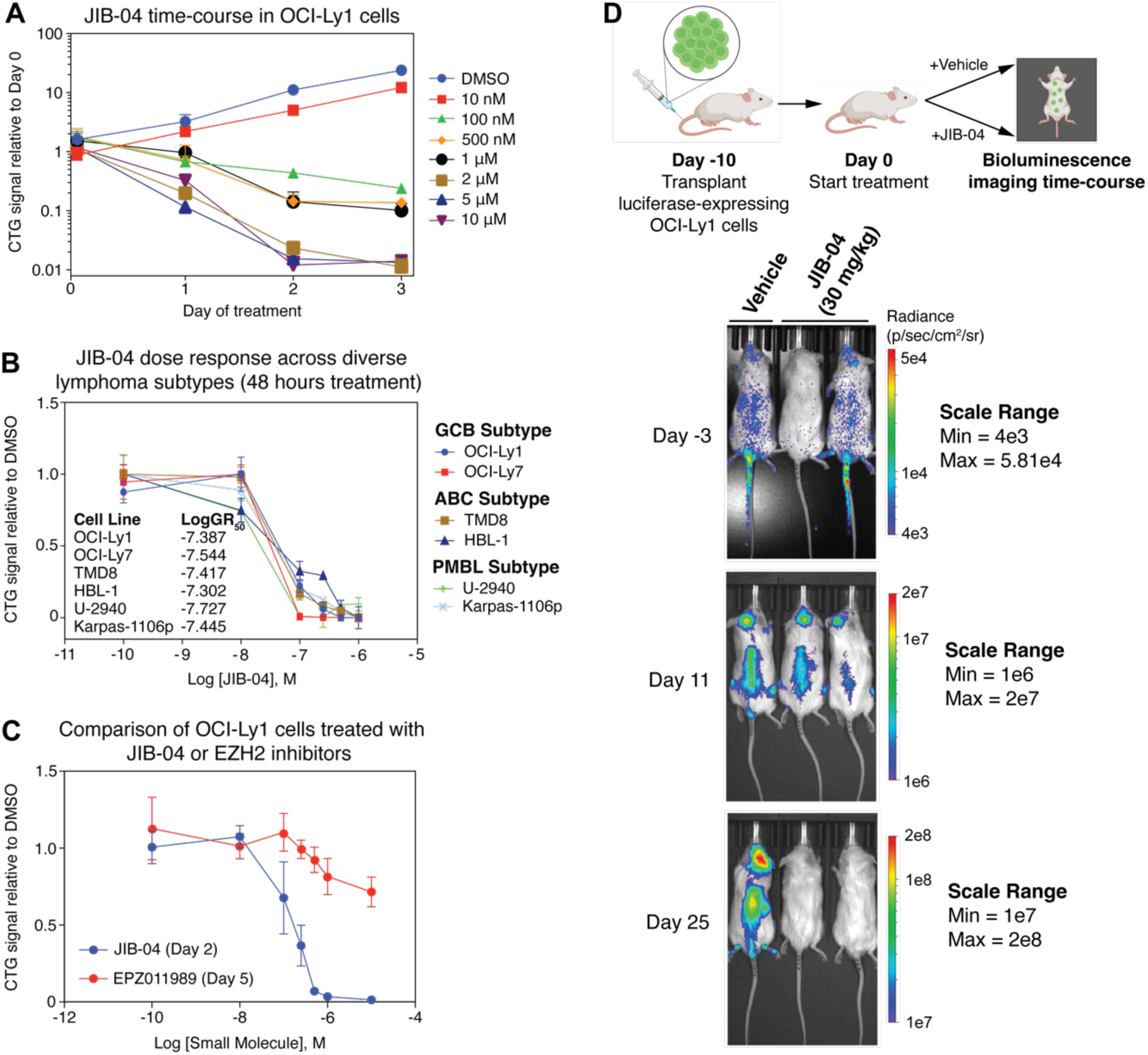
JIB-04 exhibits potent anti-cancer activity against diverse subtypes of DLBCL. **A)** CellTiter-Glo time-course of OCI-Ly1 cells treated with various doses of JIB-04 over 3 days. CTG signal at each time-point is normalized to the starting day 0 baseline signal. Error bars represent standard deviations across three biological replicates per dose and time-point. 0.1% DMSO was used as a vehicle control. **B)** JIB-04 dose titration across a broad compendium of DLBCL cell lines. CTG signal at each dose for each cell line is normalized to DMSO vehicle controls. Error bars represent standard deviations across three biological replicates per dose. **C)** Dose titration comparison of JIB-04 and EZH2 inhibitor, EPZ011989 in OCI-Ly1 cells. CTG signal at each dose is normalized to the CTG signal of 0.01% DMSO vehicle control. Error bars represent standard deviations across n=3 biological replicates per dose. **D)** *In vivo* treatment of tumor xenografts with JIB-04. Luciferase-expressing OCI-Ly1 cells were transplanted to NSG mice via tail vein injection and tumor burden was assessed after 10 days. Mice were treated daily for 25 days with JIB-04 (30 mg/kg) or DMSO vehicle control. The scale range in radiance (p/sec/cm^2^/sr) is noted for each timepoint.

We then assessed the activity of JIB-04 across a broad compendium of DLBCL cell lines. All 14 cell lines tested exhibited a 50% reduction in growth rate (GR_50_) at doses less than 250 nM (**Figure S2D-E**). The sensitivity of this broader compendium of DLBCL cells to JIB-04 treatment was observed within 48 hours (**Figure S2E**), consistent with our validation studies. Curiously, two primary mediastinal B-cell lymphoma (PMBL) cell lines, U-2940 and Karpas-1106p exhibiting genomic amplifications of the chromosome 9p locus harboring the H3K9 demethylase KDM4C were also sensitive to JIB-04 to a comparable degree as DLBCL cells (**Figure 2B**). Malignant hematopoietic cell lines of non-B cell origin were generally less sensitive to JIB-04 treatment than B cell malignancies (**Figure S2F**), suggesting a degree of B cell-specific susceptibility. JIB-04 also exhibited faster cytotoxicity kinetics than EZH2 inhibitors (**Figure 2C, S2G**), which have shown partial efficacy in EZH2-mutant GCB DLBCLs^43–45^.

Motivated by these results, we then asked whether JIB-04 could reduce tumor burden *in vivo.* We transplanted OCI-Ly1 cells bearing a constitutive luciferase reporter^47^ via tail vein injection of immunocompromised mice to generate a xenograft model of DLBCL. Daily treatment of xenografted mice with 30 mg/kg of JIB-04 over 25 days led to a substantial reduction in whole body luciferase signal in two independent treatment cohorts (**Figure 2D, S2H**). Notably, we also observed reduced intracranial tumor burden, which is a notorious site of tumor resistance to CD19-directed CAR-T cell therapies. Collectively, these data establish the *in vitro* and *in vivo* potency of JIB-04 across a broad spectrum of DLBCL subtypes.

### Heterochromatin is epigenetically rewired following JIB-04 treatment

To garner mechanistic insights into the subtype-independent sensitivity of DLBCL cells to JIB-04, we focused on understanding alterations to the epigenome and *trans-*acting chromatin regulators. Since JIB-04 is a pan-Jumonji domain inhibitor^58^, we used chromatin immunoprecipitation followed by sequencing (ChIP-seq) to profile diverse histone PTMs associated with transcriptionally active and repressed chromatin. First, we identified genomic loci with statistically significant changes in H3K4me3 and H3K27ac (**Figure S3A-B**), which are associated with transcriptionally active chromatin. The magnitude of changes was more pronounced for H3K4me3 compared to H3K27ac, where 1,819 peaks gained H3K4me3 and 1,257 peaks lost H3K4me3 following JIB-04 treatment (**Figure S3C**). Regions that lost H3K4me3 included those associated with the TLR/NF-κB arm of the BCR signaling cascade; while conversely those that gained were enriched for apoptosis, repression of Wnt/beta-catenin and the activation of inflammatory pathways (FDR-adjusted *P* value < 0.05, **Figure S3D-E**). These data suggest that JIB-04 alters chromatin to antagonize core B-cell survival programs. To further explore this, we inferred TF activity within H3K4me3 and H3K27ac ChIP-seq peaks by chromVAR^59^. Interestingly, TF motifs associated with B-cell identity, such as the IKZF and PAX families were inferred to have attenuated activity within both H3K27ac and H3K4me3 ChIP-seq peaks following JIB-04 treatment (**Figure 3A**). We validated that IKZF1, and its downstream targets IKZF3 and the BCR signaling component, SYK, were indeed downregulated at the protein level following 24 hours of JIB-04 treatment in multiple DLBCL subtypes (**Figure 3B, S3F**). Collectively, these data suggest that JIB-04 treatment is altering B-cell lineage transcriptional networks, leading to down-regulation of B-cell survival programs.

**Figure 3.**
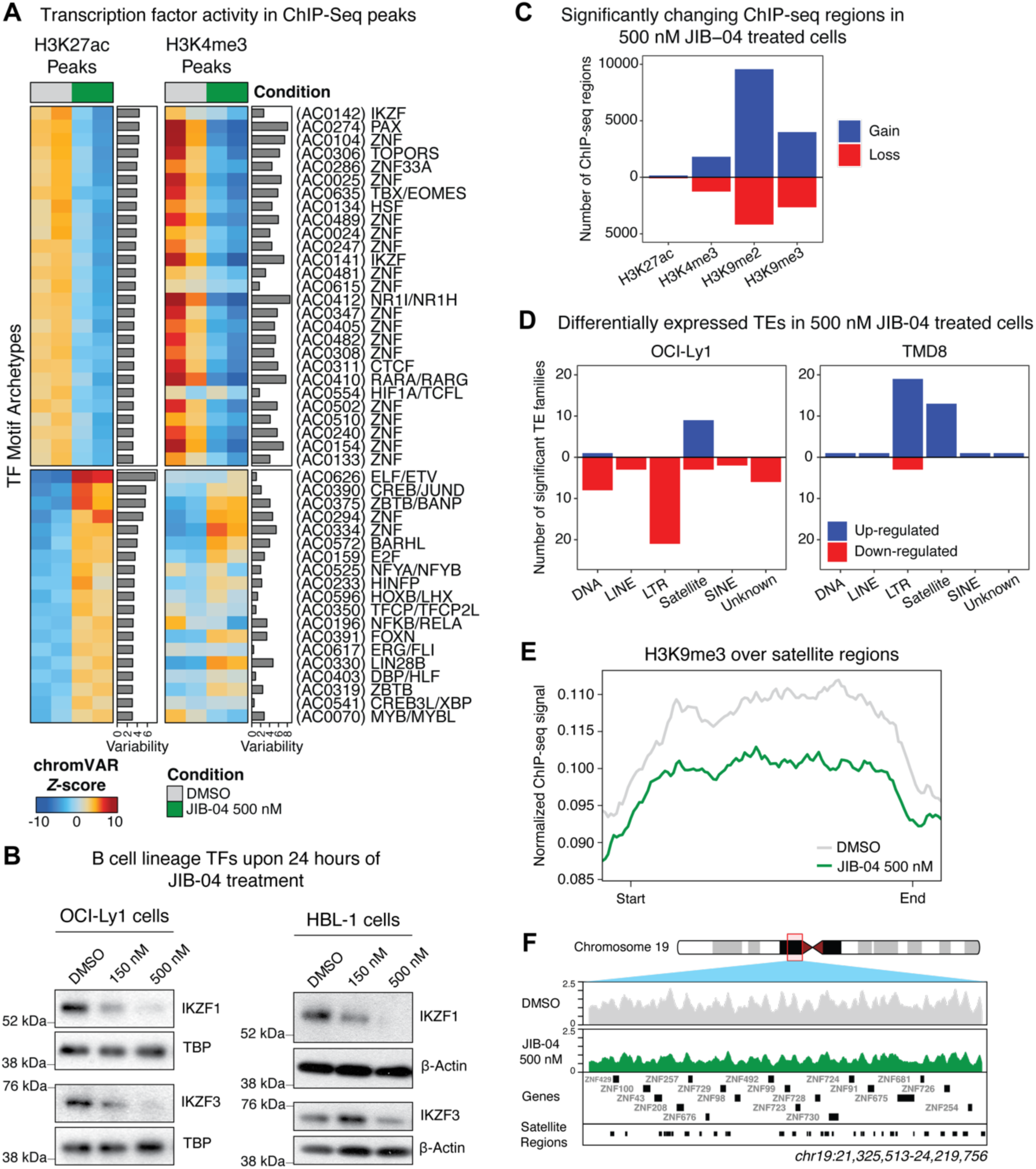
JIB-04 treatment alters the genome-wide heterochromatin landscape. **A)** Heatmap of statistically significant (FDR-adjusted *P* value < 0.05) differences in TF motif archetypes in H3K27ac and H3K4me3 ChIP-seq peaks between 500 nM JIB-04 and DMSO treated OCI-Ly1 cells. Motif archetypes are derived from Vierstra*, et al.* (2020). *Nature*. (PMID: 32728250). The archetype identifiers from the v2.1beta release are noted in parentheses and the consensus TF in the motif cluster is labeled. Color bar represents the chromVAR deviation Z-score. The standard deviation of chromVAR deviation Z-scores across all samples in a ChIP-seq experiment is plotted along the rows of the heatmaps and indicative of the variability in ChIP-seq signal over motifs between treatment conditions. **B)** Immunoblot for IKZF1 and IKZF3 from OCI-Ly1 and HBL-1 cells treated with multiple doses of JIB-04 for 24 hours. **C)** Quantification of the number of ChIP-seq regions that significantly gain or lose H3K27ac, H3K4me3, H3K9me2, or H3K9me3 in 500 nM JIB-04 versus DMSO treated OCI-Ly1 cells. ChIP-seq regions that exhibit an FDR adjusted *P-*value < 0.05 are considered statistically significant. **D)** Quantification of differentially expressed TE families between 500 nM JIB-04 and DMSO treated OCI-Ly1 cells for RepeatMasker annotated DNA, LINE, LTR, Satellite, SINE and Unknown TE classes. Differentially expressed TEs are defined as FDR-adjusted *P* value < 0.05 and log_2_(fold change) > 0.75 and log_2_(fold change) < −0.75. **E)** Normalized H3K9me3 ChIP-seq signal over satellite regions transcriptionally up-regulated in 500 nM JIB-04 versus DMSO treated OCI-Ly1 cells. **F)** Genome sequencing tracks of the 19p12 locus (chr19:21,325,513-24,219,756 in hg38 coordinates) visualizing H3K9me3 ChIP-seq in DMSO and 500 nM JIB-04 treated OCI-Ly1 cells. RepeatMasker-annotated BSR/beta satellite elements are shown in the bottom row.

Our ChIP-seq analyses also revealed global alternations with H3K4me3 and H3K27ac over multiple zinc finger (ZNF) TF motifs, where two and 17 ZNF motif families gained and attenuated activity, respectively (**Figure 3A**). These chromatin observations were consistent with 50 reproducibly differentially expressed ZNF genes after JIB-04 treatment in both subtypes of DLBCL (**Figure S3G**). Furthermore, TRIM28, the transcriptional scaffold protein that binds KRAB domain-containing ZNF proteins (KZFPs) to promote heterochromatin formation was also attenuated at the protein level in JIB-04 treated cells (**Figure S3H**). Thus, we profiled H3K9me2 and H3K9me3 histone modifications via ChIP-seq to determine how JIB-04 affects heterochromatin. Treatment of OCI-Ly1 cells with JIB-04 altered more heterochromatin loci marked by H3K9me2 (13,742 regions) or H3K9me3 (6,645 regions) compared to euchromatin loci marked by H3K4me3 (3,076 peaks) or H3K27ac (250 peaks) (**Figure 3C**). Since KZFPs principally act to target and regulate repetitive elements in the human genome, including pericentromeric heterochromatin and transposable elements, we then leveraged RNA-sequencing (RNA-seq) to quantify repeat element expression in JIB-04 treated cells. Several repetitive element families were significantly differentially expressed, particularly long terminal repeat (LTR) elements, yet we observed opposing effects in OCI-Ly1 (43 total families down-regulated, 10 up-regulated) and TMD8 cells (4 total families down-regulated, 36 up-regulated) despite both cell lines being susceptible to JIB-04 treatment (**Figure 3D**). However, expression of satellite repeat regions were consistently up-regulated in both JIB-04 treated cell lines (OCI-Ly1: 9 satellite families, TMD8: 13 satellite families, **Figure 3D**), thus we hypothesized that disruption to constitutive heterochromatin is a conserved consequence of JIB-04 treatment. Satellites were also significantly up-regulated in a dose-dependent manner across DLBCL subtypes (**Figure S3I**), further supporting this hypothesis. H3K9me3, which marks constitutive heterochromatin, was globally attenuated over differentially expressed satellite regions in a JIB-04 dependent manner (**Figure 3E**). Interestingly, the BSR/beta family of satellite elements that were transcriptionally up-regulated in JIB-04 treated cells (**Figure S3I**) were enriched in the chr19p12 locus and co-localized with a cluster of ZNF genes (**Figure 3F**). Our chromatin and transcriptional data thus suggest that JIB-04 globally alters heterochromatin, and the regulatory networks associated with B cell identity and maintenance of genome stability.

### KDM4 demethylases are targets of JIB-04 and vulnerabilities in DLBCL

We then sought to identify the direct targets of JIB-04. Motivated by our epigenomic data, we hypothesized that JIB-04 inhibits the enzymatic activity of Jumonji domain-containing H3K9 demethylases. To test this hypothesis, we first utilized transcriptional profiling datasets from DLBCL patients^60^. Among the set of H3K9 demethylases in the human genome, KDM4A and KDM4C were transcriptionally upregulated in DLBCL patients relative to healthy controls (*P* < 0.05, **Figure S4A**), suggesting potential DLBCL dependencies (**Figure S4B**). We therefore prioritized our biochemical investigations on KDM4A/C. We first used a demethylation activity assay with fully defined semi-synthetic nucleosome substrates,^61^ and identified that JIB-04 inhibited H3K9me3 demethylation by KDM4A/C proteins to a comparable degree as PDCA, a 2-OG mimic that inhibits Jumonji domain-containing demethylases (**Figure 4A-B**). KDM4A and KDM4C, but not KDM5A, were also stabilized by JIB-04 within DLBCL cells, as evidenced by cellular thermal shift assays (**Figure 4C-D, S4C**). These results support our *in vitro* biochemical evidence that JIB-04 binds and enzymatically inhibits KDM4A/C.

**Figure 4.**
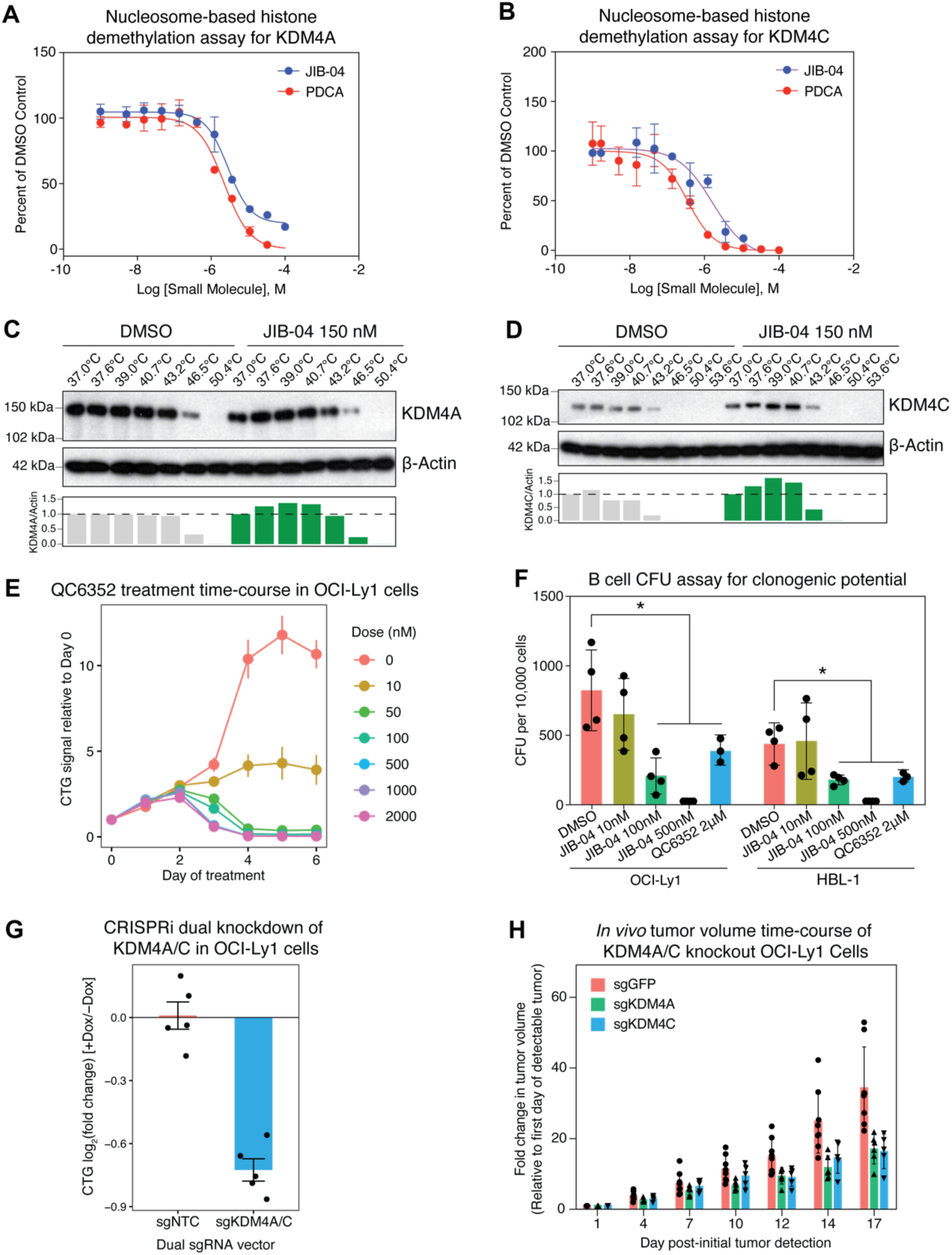
KDM4 proteins are targets of JIB-04 and vulnerabilities in DLBCL. **A-B)** KDM4A **(A)** or KDM4C **(B)** *in vitro* enzymatic activity was monitored by the conversion of H3K9me3 nucleosome substrate to H3K9me2 product across a titration of JIB-04 or PDCA (positive control). Error bars represent standard deviations across three replicates per dose. AlphaLISA luminescence signal at each dose is normalized to parallel DMSO vehicle controls. **C-D)** Cellular thermal shift assays indicating that JIB-04 stabilizes KDM4A **(C)** and KDM4C **(D)** within OCI-Ly1 cells. Expected molecular weights: β-Actin, 42 kDa, KDM4A, 150 kDa, KDM4C, 120 kDa. Relative KDM4A/C protein signal normalized to the β-Actin loading control is quantified for each sample below each blot. **E)** CellTiter-Glo time-course of OCI-Ly1 cells treated with KDM4 inhibitor QC6352 across various doses. CTG signal at each time-point is normalized to the starting day 0 baseline signal. Error bars represent standard deviations across three biological replicates per dose and time-point. 0.02% DMSO was used as a “0 nM” vehicle control. **F)** Colony forming potential of OCI-Ly1 and HBL-1 cells exposed to DMSO, JIB-04 or QC6352 within methylcellulose media. Each point represents an independent biological replicate and error bars reflect one standard deviation. Colonies were scored in a blinded manner after two weeks of culture. Asterisks indicate an FDR < 0.05, computed by comparing each drug treatment condition with DMSO controls (two-tailed Wilcoxon rank-sum test, corrected for testing of multiple modules). **G)** Fold change in CellTiter-Glo signal following 6 days of CRISPRi-mediated dual knockdown of KDM4A/C in comparison to non-targeting control sgRNAs. CRISPRi machinery is dox inducible and thus fold changes for each sgRNA condition are relative to -dox controls. Each point represents an independent biological replicate.**H)** *In vivo* tumor volume time-course of tumor xenografts in NSG mice. Cas9-expressing OCI-Ly1 cells that received sgRNAs to knock out KDM4A, KDM4C or GFP (non-targeting control) were suspended in matrigel and transplanted subcutaneously to NSG mice. Tumor volumes were measured over 17 days, with measurements relative to the first day of detectable tumor. Each point represents relative tumor volumes from independent mice.

We then utilized another KDM4 inhibitor QC6352^62^ (which was not represented in our original phenotypic screens), to further investigate KDM4-specific targeting in DLBCL. QC6352 phenocopied the cytotoxicity elicited by JIB-04, albeit with differing kinetics (**Figure 4E, S4D**). Treatment of OCI-Ly1 and HBL-1 cells with either JIB-04 or QC6352 reduced colony forming units in a long-term clonogenic assay (*P* < 0.05, **Figure 4F**), further suggesting that KDM4A/C are needed for the subtype-independent survival of DLBCL cells. To directly perturb KDM4A/C and to exclude the possibility of off-target drug effects contributing to cellular cytotoxicity, we performed multiplexed CRISPRi knockdown of KDM4A/C in OCI-Ly1 cells (**Supplemental Table S2**). We derived multiple, independent clones that harbored doxycycline-inducible CRISPR machinery, and unexpectedly the clones exhibited varying baselines of KDM4A/C protein expression (**Figure S4E**). However, robust knockdown of KDM4A/C protein led to a significant decrease in cell viability across all clones over six days (*P* < 0.05, **Figure 4G, S4E-F**). This KDM4A/C dependence was also corroborated by a CRISPR/Cas9-mediated knockout approach (**Figure S4G-H**). Finally, knockout of KDM4A/C also reduced tumor burden within xenograft models of DLBCL *in vivo* (**Figure 4H**). Our chemical and genetic evidence suggest that KDM4A/C are subtype-agnostic vulnerabilities in DLBCL.

### Modulation of KDM4 induces cell-intrinsic inflammation via replication stress

We were then motivated to understand how inhibition of KDM4A/C results in cytotoxicity in DLBCL cells. We utilized whole-transcriptome profiling of JIB-04 treated OCI-Ly1 and TMD8 cells and observed that 90% of the transcriptional variance in the data (deduced by principal component analysis) was treatment driven (**Figure S5A**). Given the global transcriptional changes induced by JIB-04, we then identified differentially expressed genes and subsequently performed gene set enrichment analysis (GSEA) using Hallmark gene sets^63^. Notably, induction of the p53 pathway and inflammatory signaling via NFkB, as well as suppression of MYC target genes, were enriched in GCB and ABC subtypes (FDR-adjusted *P* value < 0.05, **Figure 5A-B**). Specifically, *ATF3, TP63,* and *CDKN1A* were consistently upregulated across DLBCL both subtypes, as well as induction of an interferon signature (*IRF1, IRF7, IRF9, STAT5A, ISG20, IFIT2*), suggestive of DNA damage-induced inflammation (**Figure 5C**). In support of this possibility, we also observed down-regulation of *ZNF587* (**Figure 5C**), which was recently implicated in safeguarding DLBCL cells against genome instability during DNA replication^64^.

**Figure 5.**
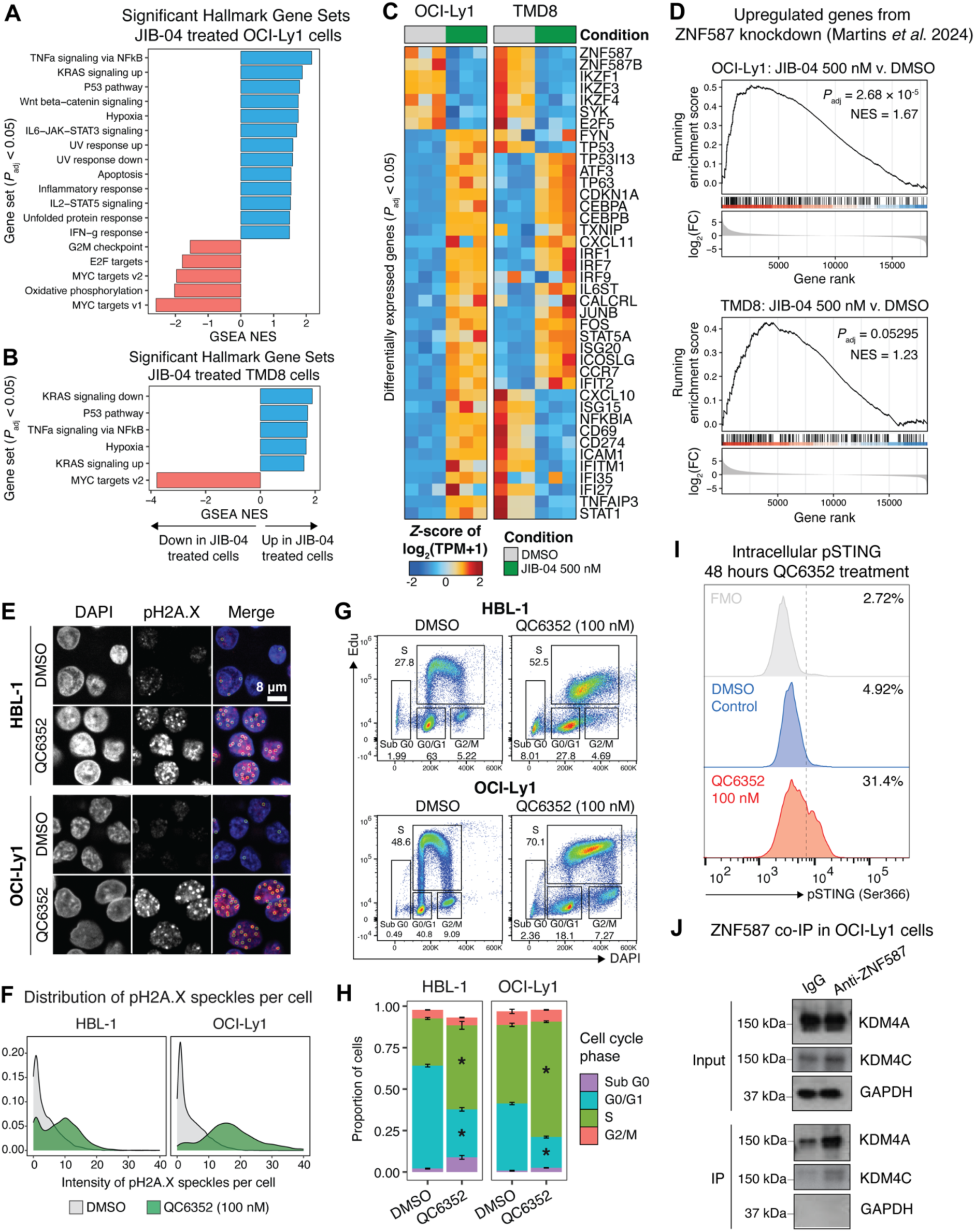
Modulation of KDM4 induces cell-intrinsic inflammation via replication stress. **A-B)** Gene set enrichment analysis (GSEA) of Hallmark gene sets on JIB-04 treated OCI-Ly1 **(A)** and TMD8 **(B)** cells. All terms depicted are statistically significant following correction for multiple hypothesis testing (FDR adjusted *P-*value < 0.05). Positive NES values reflect gene sets significantly enriched within JIB-04 treated cells, whereas negative NES values reflect gene sets significantly enriched in DMSO treated cells. **C)** Heatmap of statistically significant (FDR adjusted *P-*value < 0.05) differentially expressed genes between 500 nM JIB-04 and DMSO treated cells. Each column represents independent biological replicates. log_2_(TPM+1) expression values are Z-score standardized along rows. **D)** GSEA on genes upregulated following ZNF587 knockdown in DLBCL cell lines from Martins, *et al.* 2024 (PMID: 38345497). Genes were ranked based on descending DESeq2 log_2_(fold change) between JIB-04 and DMSO treated cells. Positive NES scores reflect gene set enrichment within 500 nM JIB-04 treated cells compared to DMSO treated cells. GSEA *P-*values were FDR-corrected for multiple hypothesis testing. **E)** Representative microscopy of HBL-1 and OCI-Ly1 cells treated with either 0.02% DMSO or 100 nM QC6352 for 48 hours and stained for DAPI and pH2A.X (Ser139). Detection of computationally inferred pH2A.X speckles is visualized in the merged images. Scale bar: 8 µm. **F)** Distribution of pH2A.X speckle intensity from 0.02% DMSO or 100 nM QC6352 treated OCI-Ly1 and HBL-1 cells. **G)** Flow cytometry quantification of replication (EdU incorporation intensity) and DNA content (DAPI) in HBL-1 and OCI-Ly1 cells after treatment with 0.02% DMSO or 100 nM QC6352 for 48 hours. Cell cycle phases are annotated within plots. **H)** Quantification of the proportion of cells in each cell cycle phase from **(G)** across three biological replicates. Error bars represent standard deviations. Asterisks indicate *P* < 0.05 based on Student’s *t-*test for QC6352 versus DMSO. **I)** Flow cytometry for intracellular pSTING (Ser366) after treatment with DMSO or 100 nM QC6352 for 48 hours. **J)** ZNF587 immunoprecipitates (IPs) with KDM4A and KDM4C demethylases. Total protein extract from OCI-Ly1 cells (top set of blots, “Input”) was subject to IP with anti-ZNF587 or IgG isotype control followed by immunoblotting (bottom set of blots, “IP”).

We thus hypothesized that inhibition of KDM4A/C elicits cytotoxicity through DNA damage. Utilizing transcriptional data from Martins *et al*.^64^, we created a gene set comprising genes that were consistently up-regulated upon knockdown of *ZNF587* across various DLBCL cell lines (**Supplemental Table S3**). This *ZNF587* knockdown gene signature was significantly enriched within JIB-04 treated cells across various doses and in both DLBCL subtypes (FDR-adjusted *P* value ≤ 0.05, **Figure 5D, S5B**), suggesting that the genome instability induced by knockdown of *ZNF587* is also reflected in JIB-04 treated cells. These results were corroborated by evidence of DNA damage via increased Ser-139 H2A.X phosphorylation (γH2A.X) in KDM4 inhibitor-treated OCI-Ly1 and HBL-1 cells (*P* < 0.001, **Figure 5E-F**). Curiously, all S phase cell cycle genes were transcriptionally dysregulated in JIB-04 compared to DMSO treated cells (**Figure S5C**). These observations coupled with prior literature indicating heterochromatin regions are late-replicating in the cell cycle^65–67^ prompted us to then assess DNA replication through the incorporation of EdU. Following 48 hours of treatment with KDM4 inhibitors, significantly more cells were in S phase (DMSO: 47.4%, 100 nM QC6352: 69.6%, *P* < 0.001) and correspondingly fewer cells in G0/G1 (DMSO: 40.5%, 100 nM QC6352: 18.6%, *P* < 0.001) (**Figure 5G-H**). These data implicated cell cycle arrest through replication stress. Cells treated with KDM4 inhibitors also exhibited activation of the STING pathway (Ser-366 phosphorylation) (**Figure 5I, S5D**), and treatment with STING inhibitor, H-151 partially rescued KDM4 inhibitor-induced cytotoxicity (**Figure S5E**).

Finally, since inhibition of KDM4A/C molecularly phenocopied knockdown of *ZNF587*, we posited that both sets of proteins may cooperate to protect DLBCL cells from replication stress. Indeed, co-immunoprecipitation of ZNF587 and KDM4A/C from OCI-Ly1 cells identified a physical association between the proteins (**Figure 5J**). Thus, our data suggests that KDM4 proteins interact with KZFPs to protect from genome instability, and inhibiting KDM4 demethylases results in DNA damage and replication stress, instigating inflammation through cGAS-STING pathway activation.

### KDM4 inhibition is cytotoxic for various human cancers

Interrogation of TCGA data indicated that KDM4A/C are frequently differentially expressed in human cancers (KDM4A: 70%, 14/20, TCGA cancer types; KDM4C: 60%, 12/20 TCGA cancer types, **Figure S6A**), so we next asked if modulating the Jumonji demethylases could therapeutically extend beyond lymphoma. In this regard, KDM4A/C expression was significantly associated with overall survival in the TCGA COAD patient cohort (*P* = 0.0049), and HCT116 colorectal cancer cells were susceptible to KDM4 inhibitors, with the cytotoxicity partially rescued by cGAS-STING inhibition (**Figure S6B-C**). These data further support the generality of the molecular mechanism of KDM4-mediated cytotoxicity that we identified in DLBCL, and suggest that KDM4 inhibition could be a therapeutic vulnerability in other malignancies.

### High-throughput small molecule screens identify novel KDM4 inhibitors

While the dysregulation of KDM4 family members across human cancers warrants further therapeutic investigation, current tool compound inhibitors are likely unsuitable for clinical development. We thus developed a high-throughput screening workflow to identify novel KDM4 demethylation inhibitors that could represent a path to therapeutics (**Figure 6A**). We designed a primary screen utilizing a H3K9me3 histone peptide substrate and recombinant KDM4C enzyme, with relative demethylation activity monitored on the AlphaLISA platform by detection of the H3K9me1 product (**Figure 6A**). After confirming reproducibility and dynamic range of the demethylation assay for high-throughput screening (Z’ = 0.701, **Figure S7A**), we screened a library of 50,000 small molecules from ChemBridge. The primary screen was highly reproducible across screening replicates (Pearson *r* = 0.98, **Figure S7B**), with 961 molecules exhibiting a *Z-* score < −3, constituting a 1.9% hit rate (**Figure 6B, Supplemental Table S4**). Hits were filtered to exclude pan-assay inhibitors and selectively curated to encompass a chemically diverse set of molecules for further study (see Methods). We rescreened a prioritized set of 297 molecules in the same histone peptide assay, and counter screened with a H3K9me3 semi-synthetic nucleosome substrate to further support that the molecules are enzymatic inhibitors (**Figure S7C-D**). Of the 297 molecules, 252 (84.8%) reconfirmed in the histone peptide assay, with 177 hits (59.5%) in the nucleosome assay, and 159 hits (53.5%) in both substrate formats (**Figure 6C, Supplemental Table S5**). In parallel we performed an AlphaLISA TruHits assay on the 297 prioritized molecules to exclude false positive hits that interfere with the AlphaLISA demethylation assay, which identified 89 molecules (29.9%) that qualified as true hits (**Figure 6D**). Across all our secondary screens, 56 molecules were true hits in both nucleosome and peptide assays (18.8% of prioritized primary screen hits, **Figure 6E, Supplemental Table S5**). Using this high confidence set of small molecules, we selected the top 20 based on nucleosome demethylation activity and performed dose titration experiments on nucleosome substrates (**Figure S7E**). Here, four molecules exhibited potent dose responses with IC_50_ values < 10 µM (**Figure 6F**). Structurally similar compounds to these four showed similar inhibition (**Figure S7F**), suggesting conserved elements central for inhibitory function.

**Figure 6.**
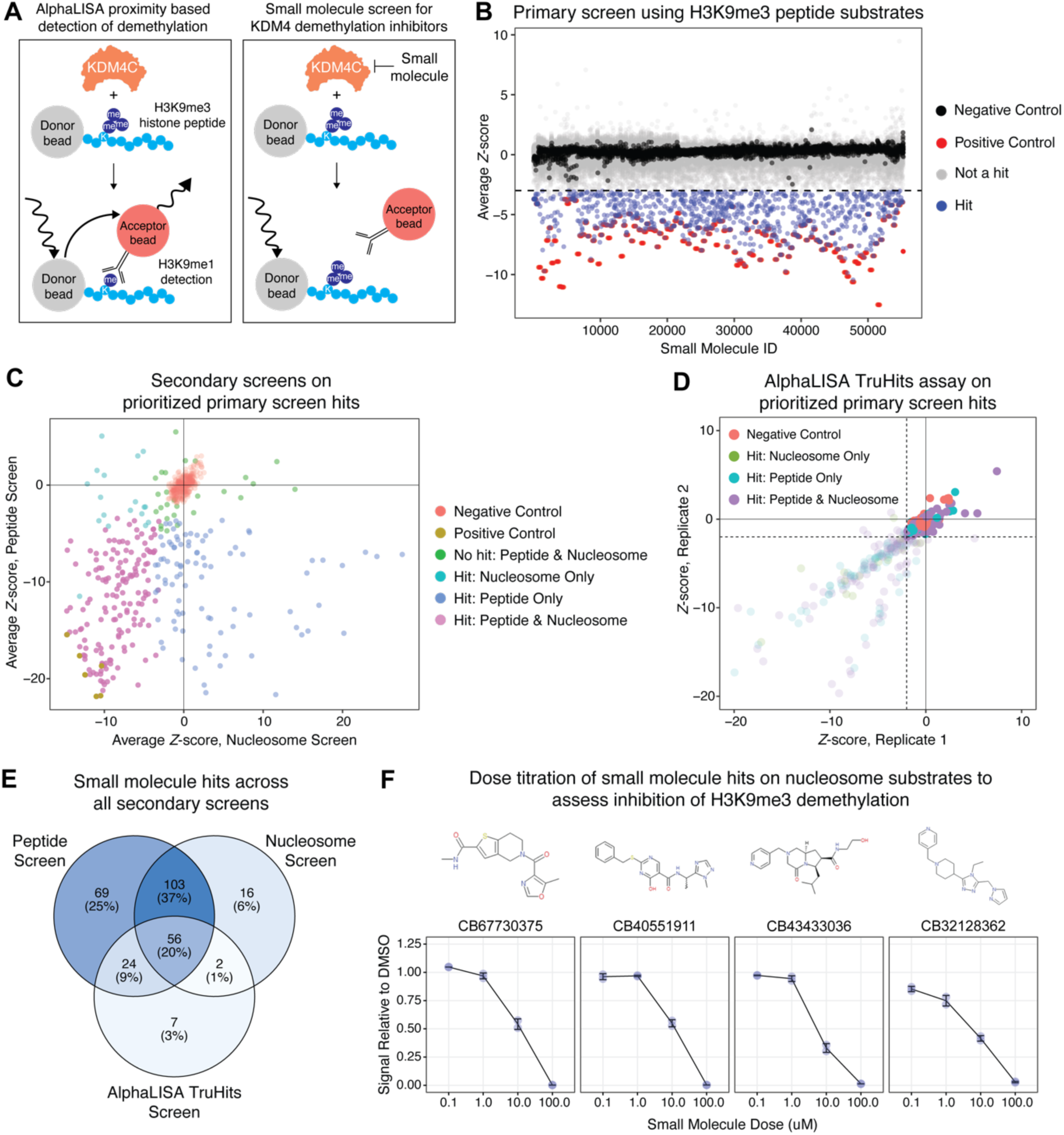
High-throughput small molecule screens identify novel KDM4 inhibitors. **A)** Experimental schematic to identify novel KDM4 inhibitors by high-throughput small molecule screening. Left panel: Recombinant human KDM4C enzyme is incubated with biotinylated H3K9me3 histone peptide substrate; followed by sequential addition of anti-H3K9me1 and streptavidin donor / protein A acceptor beads. Excitation (680 nm) of donor beads leads to emission (615 nm) from proximal acceptor beads, providing a quantitative output to what degree all reactants are bridged by H3K9me1 enzyme product. Right panel: we then leveraged this *in vitro* system to screen libraries of small molecules to identify inhibitors of H3K9me3 demethylation. Small molecule ligands that inhibit the demethylation of recombinant KDM4C would attenuate the AlphaLISA signal relative to DMSO vehicle controls. **B)** Primary screen to identify small molecule inhibitors of KDM4C enzymatic activity on histone peptide substrates. The ChemBridge 2020 library (50,000 small molecules) was screened in an arrayed manner as in **(A)**. Replicate reproducibility and relative signal of negative (DMSO) and positive (CPI-455) control reactions were used to confirm assay robustness and derive a hit-threshold. Screens were performed in duplicate and small molecules with a Z-score < −3 in both reactions defined as hits. Visualized is the average Z-score across both screening replicates. **C)** 297 small molecule hits from the primary screen in **(B)** were rescreened with KDM4C and H3K9me3 histone peptide or semi-synthetic nucleosome substrates in duplicate. Small molecules with a Z-score < −3 in both screening replicates and across both substrate classes were of greatest interest (subject to TruHits filtering). **D)** 297 small molecule hits from the primary screen in **(B)** were rescreened with an AlphaLISA TruHits assay to filter false-positive hits that broadly interfere with the format. Candidates with an TruHits value less than two standard deviations of the mean of DMSO controls (transparent points) were considered false-positives and removed from further investigation. **E)** Venn diagram of small molecule hits across all secondary screens, resulting in 56 high confidence candidates. **F)** Dose titration of representative high confidence candidates (chemical structure noted above each plot) on H3K9me3 nucleosome substrates. AlphaLISA demethylation signal is normalized to DMSO controls. Error bars represent standard deviations across replicates.

We then leveraged recent AI models for predicting protein-ligand binding to further explore a representative small molecule: CB43433036, which exhibited the lowest IC_50_ on nucleosome substrates (IC_50_ = 6.84 µM, **Figure 6F**). AI-Bind^68^ was used to predict putative binding sites on KDM4C and reassuringly implicated the Jumonji domain: the enzymatic active pocket that coordinates Fe(III)- and alpha-ketoglutarate-dependent histone demethylation (**Figure 7A**). Molecular docking simulations corroborated this prediction, suggesting that CB43433036 forms polar bonds with residues Y134, H190, E192, H278, K569, and R572 of KDM4C (**Figure 7B**). Unexpectedly, the model also predicted a binding site in a PHD domain of KDM4C, which recognizes methylated histone 3 tails. The small molecule ligand was predicted to dock within a pocket in the PHD domain and form polar bonds with G731, E736, and Q795 (**Figure 7C**). Collectively, our screens identified small molecule ligands that inhibit KDM4 demethylases and can potentially serve as useful scaffolds for further drug development endeavors.

**Figure 7.**
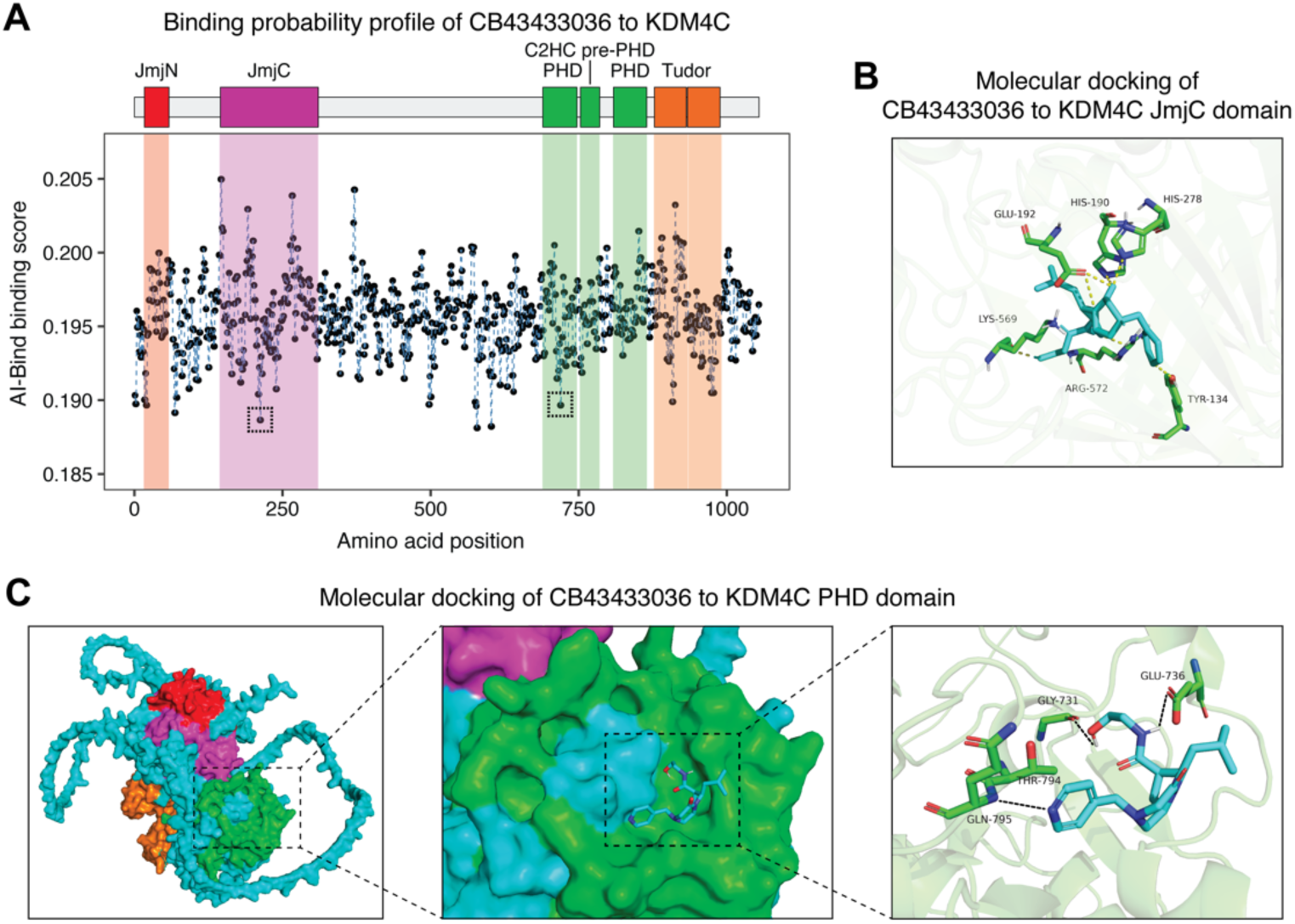
Molecular docking simulations of small molecule ligands with KDM4C. **A)** Binding probability profile from AI-Bind^68^ for small molecule ligand CB43433036 and human KDM4C (UniProt: Q9H3R0). Amino acid trigrams were perturbed throughout the KDM4C amino acid sequence to determine any influence on the binding prediction. Valleys in the binding profile (marked by dashed boxes) are indicative of putative binding sites. Top: annotation of UniProt protein domains along the KDM4C amino acid sequence. **B)** Three-dimensional structure of small molecule ligand CB43433036 in complex with the KDM4C Jumonji domain determined through molecular docking simulations and visualized in PyMOL. Dashed arrows indicate molecular interactions between the small molecule ligand and KDM4C amino acids. **C)** Molecular docking simulations of CB43433036 and the KDM4C PHD domains. Left: AlphaFold3 predicted structure of human KDM4C. Protein domains are colored as in **(A)**. Middle: CB43433036 in complex with the KDM4C PHD domain determined from AutoDock Vina. Right: Visualization of polar molecular contacts between CB43433036 and KDM4C amino acids.

## DISCUSSION

We identified H3K9 demethylases, particularly the KDM4 family, as subtype-agnostic vulnerabilities in DLBCL and PMBL. We determined that inhibition of KDM4 demethylases leads to epigenetic rewiring of heterochromatin, antagonizing transcriptional networks central for B-cell identity and eliciting DNA replication stress. The cytotoxicity induced by KDM4A/C modulation was also observed in colorectal cancer cells, suggesting the KDM4 family as an anti-cancer target in several human malignancies. Our work also advances therapeutic efforts to target cancer vulnerabilities by a stringent screening approach with counterscreens to a physiological enzyme substrate, thus increasing confidence of potential inhibitor candidates.

The KDM4 family, particularly KDM4C, has previously been implicated in cell proliferation and cell cycle control^69–73^. Our data reveal that KDM4 demethylases are critical for faithful maintenance of heterochromatin and cell cycle competence, since chemical or genetic disruption of the family attenuated proliferation and lead to cytotoxicity through DNA replication stress. These findings are consistent with reported roles of KDM4 demethylases in regulating mitotic chromosome segregation and genome stability^74–77^. Notably, DLBCL cells are prone to replication stress^78^, particularly stalling and collapse of replication forks, potentially suggesting why DLBCL cells are susceptible to KDM4 inhibition. Furthermore, we observed that significantly more genomic loci gain rather than lose H3K9me2 or H3K9me3 in JIB-04 compared to DMSO treated cells. This global alteration in histone methylation is consistent with the role for KDM4A in antagonizing HP1γ to control chromatin accessibility during DNA replication^79^. Finally, our observation of cGAS-STING pathway activation in KDM4A/C inhibited cells reinforces the notion of DNA damage-induced cell-intrinsic inflammation, and is consistent with prior reports^76^. However, unlike prior reports^76^, we did not observe changes to immune checkpoint markers in KDM4 inhibited OCI-Ly1 and TMD8 cells, similar to observations of ZNF587 knock-down in OCI-Ly1 and U2932 cells^64^. This discrepancy may be inherent to DLBCL and the limited overall efficacy of immune checkpoint blockade in DLBCL^80^.

Our data revealed an unexpected finding that the activity of several zinc finger proteins was globally altered in KDM4A/C inhibited cells. KZFPs specifically contribute to heterochromatin maintenance at repetitive elements in the human genome and recently, expression of a cluster of KZFPs have been associated with poor prognosis in DLBCL patients^64^. Our demonstration that KDM4 demethylases co-immunoprecipitate with ZNF587 mechanistically reconciles the KZFP-driven regulatory network associated with poor prognosis^64^, and the function of KDM4 demethylases in genome integrity^74–76,79^. Our data also imply that targeting KDM4 demethylases could be a therapeutic approach to undermine the KZFP regulatory network associated with poor prognosis in DLBCL patients. The physical association between both sets of proteins raises the possibility that KZFPs direct the localization of KDM4 demethylases in the genome, afforded through their DNA binding domains, although further investigation is needed to clarify the extent of this molecular cooperation.

The dysregulation of KDM4 functions in multiple human cancers motivates the need for potent inhibitors as probes of biochemical function and as potential therapeutics. Significant effort has been devoted toward developing KDM4 inhibitors with mixed success^62,81–83^. Developing inhibitors to chromatin factors is challenging in general for numerous reasons: multiple orthologues bearing remarkable structural similarity, having overlapping recruitment and activity manifesting in compensation, discrepancies between *in vitro* and cellular assays owing to differences in available substrates, and incomplete delineation of molecular mechanisms. Epigenetic regulators operate in the context of nucleosomes, but their inhibitors are typically identified by high throughput screens on reductive histone peptide substrates. An example of this disconnect comes from the SETD8 inhibitor, UNC0379, which is a potent *in vitro* tool compound (IC_50_ ∼ 7.0 µM), but effectively lacking any cellular activity owing to SETD8-nucleosome association causing a conformational change in the enzyme active site that UNC0379 cannot compete against^84^. In our small molecule screens for KDM4 inhibitors we designed our hit identification campaign to include both histone peptides and nucleosomes as substrates and selected high confidence compounds which inhibited KDM4C-mediated lysine demethylation on both substrates. Indeed, the top four molecules from our validation screens exhibited potent (IC_50_ ≤ 10 µM), dose-dependent activity on nucleosome substrates. We performed further *in silico* validation of these molecules using AI-guided protein-ligand binding predictions and unexpectedly identified that one small molecule hit was predicted to bind the PHD domain of KDM4C. While the function of many PHD-fingers in Jumonji domain-containing lysine demethylases remain unclear, this finding potentially allows for chemical probes to dissect their utility in KDM4 proteins and to target PHD domains more generally in other chromatin modifying enzymes. Overall, we anticipate that the chemical series discovered as part of this study could serve as scaffolds for more potent KDM4 inhibitors, and targeting of KDM4 demethylases could potentially advance therapeutic approaches for lymphoma and other types of cancer.

## ACKNOWLEDGMENTS

We thank Sweta Mishra, Brian Strahl, and members of the Daley and Blainey labs for invaluable scientific advice. We thank Mark Namchuk and Paul Cherfison for technical advice on small molecule screening and prioritization of small molecule candidates. MN was supported by a National Science Foundation Graduate Research Fellowship. DKJ was supported by sub-award number 101330A from the National Heart, Lung and Blood Institute (U01HL099997-07). GQD acknowledges funding from the Leukemia Lymphoma Society, Translational Research Program Award (TRP-6546-18). GQD and TEN acknowledge support from the National Heart, Lung and Blood Institute (U01HL134812-04), and National Institute of Diabetes and Digestive and Kidney Disease (NIDDK-R24DK092760). TEN also acknowledges the support of NIDDK-1R01DK098241. HL is supported by NIH (AG056318, AG61796, CA208517 and CA229100), the Glenn Foundation for Medical Research, Mayo Clinic Center for Biomedical Discovery, Center for Individualized Medicine, Mayo Clinic Cancer Center, and the David F. and Margaret T. Grohne Cancer Immunology and Immunotherapy Program. Epicypher acknowledges grants 2R44GM116584 and 2R44GM117683. PCB acknowledges funding from an NIH New Innovator Award (1DP2HL141005). YS is an American Cancer Society Research Professor and is supported in part by the National Cancer Institute Outstanding Investigator Award (R35 CA210104).

## AUTHOR CONTRIBUTIONS

MN and DKJ designed and executed experiments. BL performed ChIP-seq. CK performed flow cytometry and CFU assays. AM performed the small molecule screen for novel KDM4 inhibitors. CL assisted with Edu cell cycle assays. VM assisted with animal experiments. VM and LH performed immunoblots. AT performed co-IPs. MM performed RNA-seq library construction. MRM, AV, ZBG, and SAH performed the histone demethylase assay with guidance from MCK. TMS assisted with *γ*H2A.X microscopy and image analysis. MN, DKJ, CZ and YQ performed computational analysis. BC and MAS provided resources and analyzed data. TEN, DD, MCK, TMS, YS, HL, MMS, PCB, and GQD supervised the research and provided funding. MN prepared the figures and wrote the manuscript with GQD and with input from all authors.

## SUPPLEMENTAL FIGURES

**Supplemental Figure 1.**
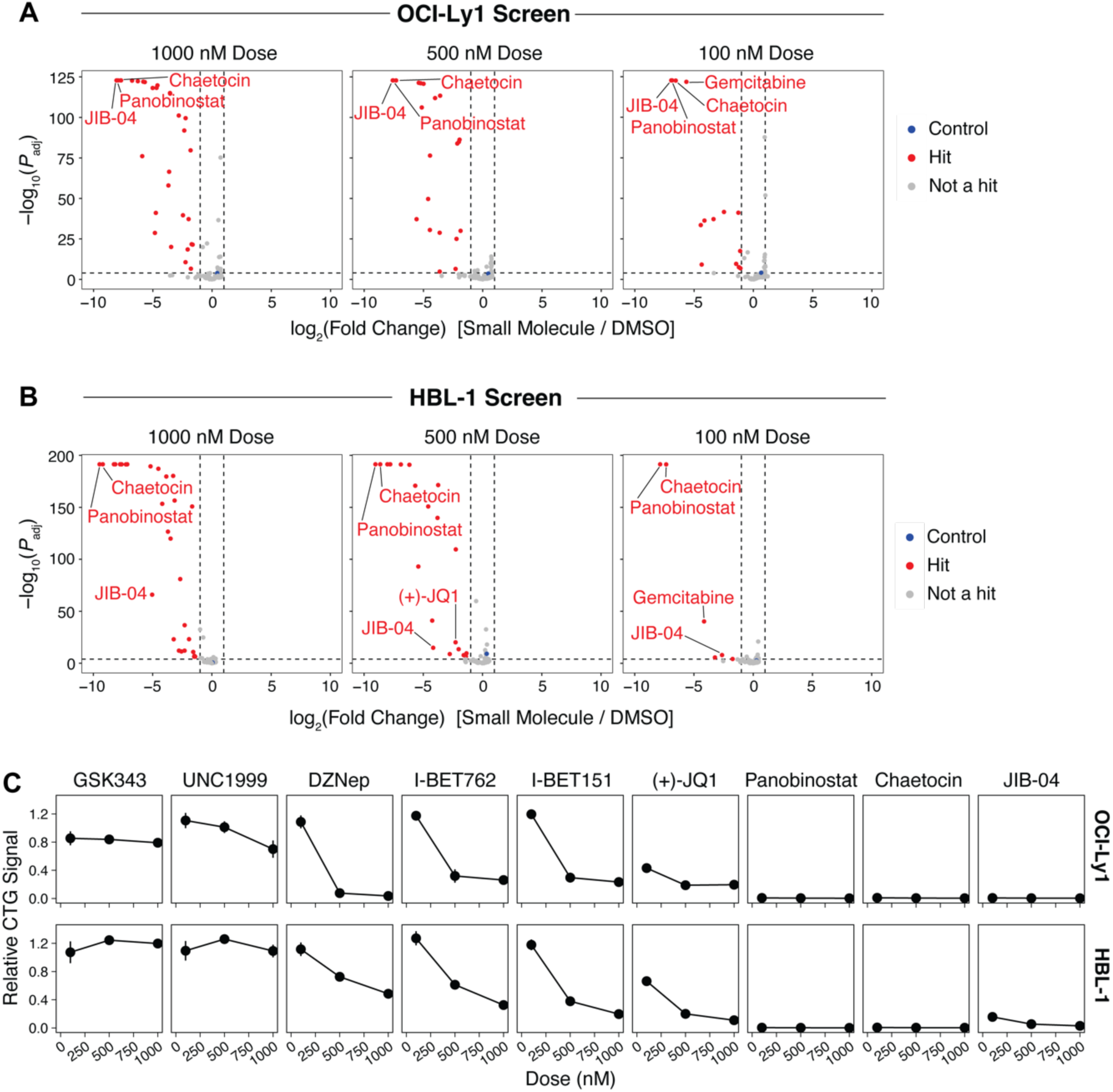
**A-B)** Volcano plots of compounds from the Cayman Chemicals Epigenetics Library assayed at 1 μM, 500 nM and 100 nM in OCI-Ly1 **(A)** and HBL-1 **(B)** cells. Fold changes are of CellTiter-Glo luminescence values after five days treatment relative to 0.01% DMSO controls. Small molecules exhibiting a log_2_(fold change) < −1 and FDR-adjusted *P* value < 0.0001 were defined as hits. **C)** Individual growth curves after chemical inhibition of EZH2 (GSK343, UNC1999, 3-Deazaneplanocin A), BET bromodomains (I-BET762, I-BET151, (+)-JQ1), HDACs (Panobinostat), SUV39H1/2 (Chaetocin) or Jumonji domains (JIB-04) from the OCI-Ly1 and HBL-1 screens after five days treatment. Presented CellTiter-Glo signal per dose is relative to DMSO controls. Each point reflects the average across three replicates and error bars represent standard deviation.

**Supplemental Figure 2.**
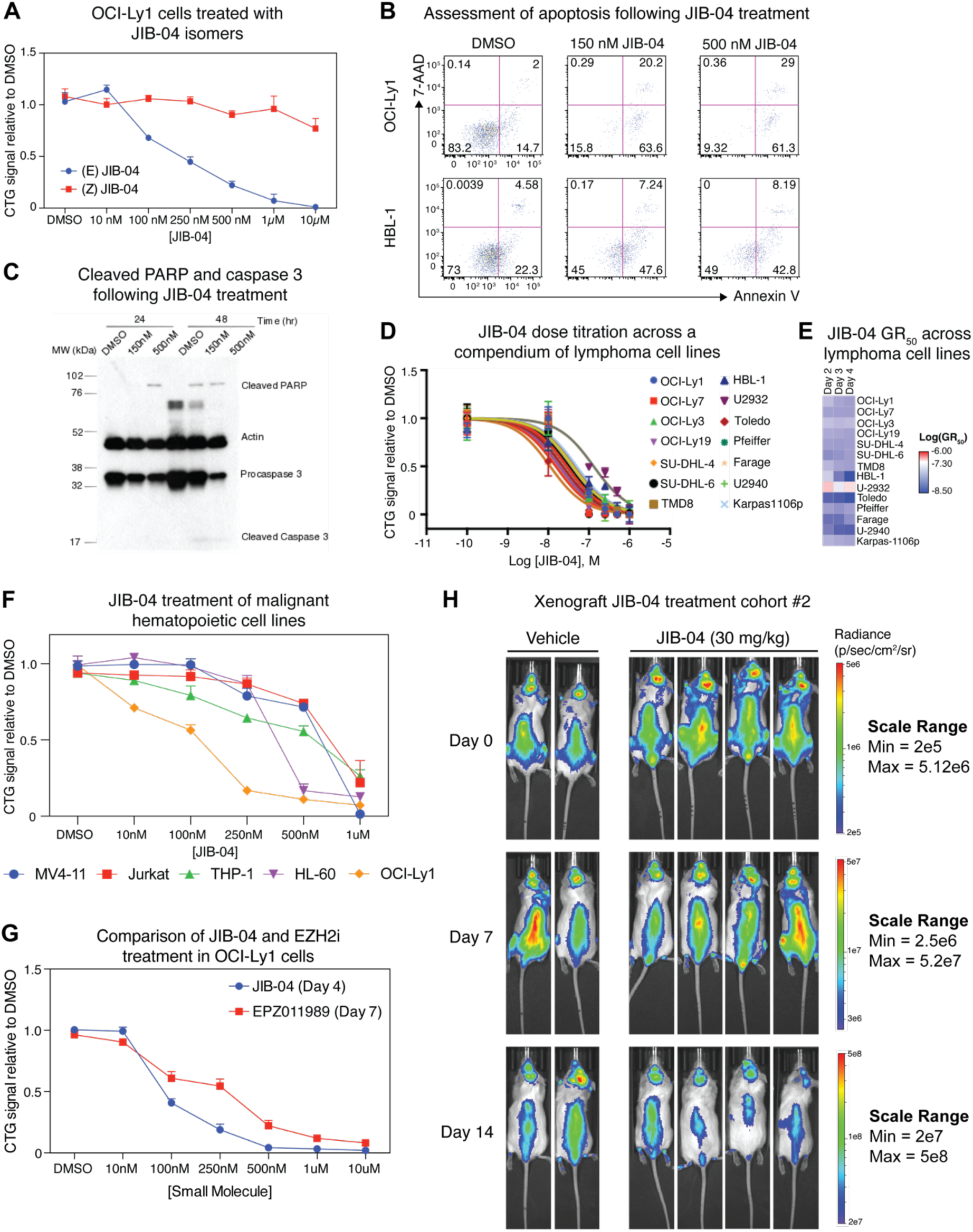
**A)** Dose titration of JIB-04 (*E*) and (*Z*) isomers on OCI-Ly1 cells. CellTiter-Glo luminescence at each dose after 48 hours is normalized to 0.1% DMSO vehicle controls. Error bars represent standard deviation over three biological replicates. B) Flow cytometry quantification of apoptosis via Annexin V and 7-AAD staining of OCI-Ly1 and HBL-1 cells treated with JIB-04 for 48 hours. C) Western-blot for cleaved PARP and cleaved caspase 3 from OCI-Ly1 treated with DMSO, 150 nM JIB-04, and 500 nM JIB-04 after 24 hours and 48 hours. D) Dose titration of JIB-04 on a broad compendium of lymphoma cell lines after 48 hours of treatment. CellTiter-Glo luminescence values at each dose are normalized to DMSO controls within each cell line. Error bars represent standard deviation over three biological replicates. E) Quantification of GR_50_ values for each cell line treated with JIB-04 from **(D)**. F) Dose titration of JIB-04 on a diverse set of malignant hematopoietic cell lines after 48 hours of treatment. CellTiter-Glo luminescence values at each dose are normalized to DMSO controls within each cell line. Error bars represent standard deviation over three biological replicates. G) Comparison of OCI-Ly1 cells after treatment with JIB-04 and EPZ011989 (EZH2 inhibitor) for four or seven days, respectively. CellTiter-Glo luminescence values at each dose are normalized to DMSO controls. Error bars represent standard deviation over three biological replicates. H) Treatment of NSG mice bearing tumor xenografts with 30 mg/kg JIB-04, performed as in Figure 2D.

**Supplemental Figure 3.**
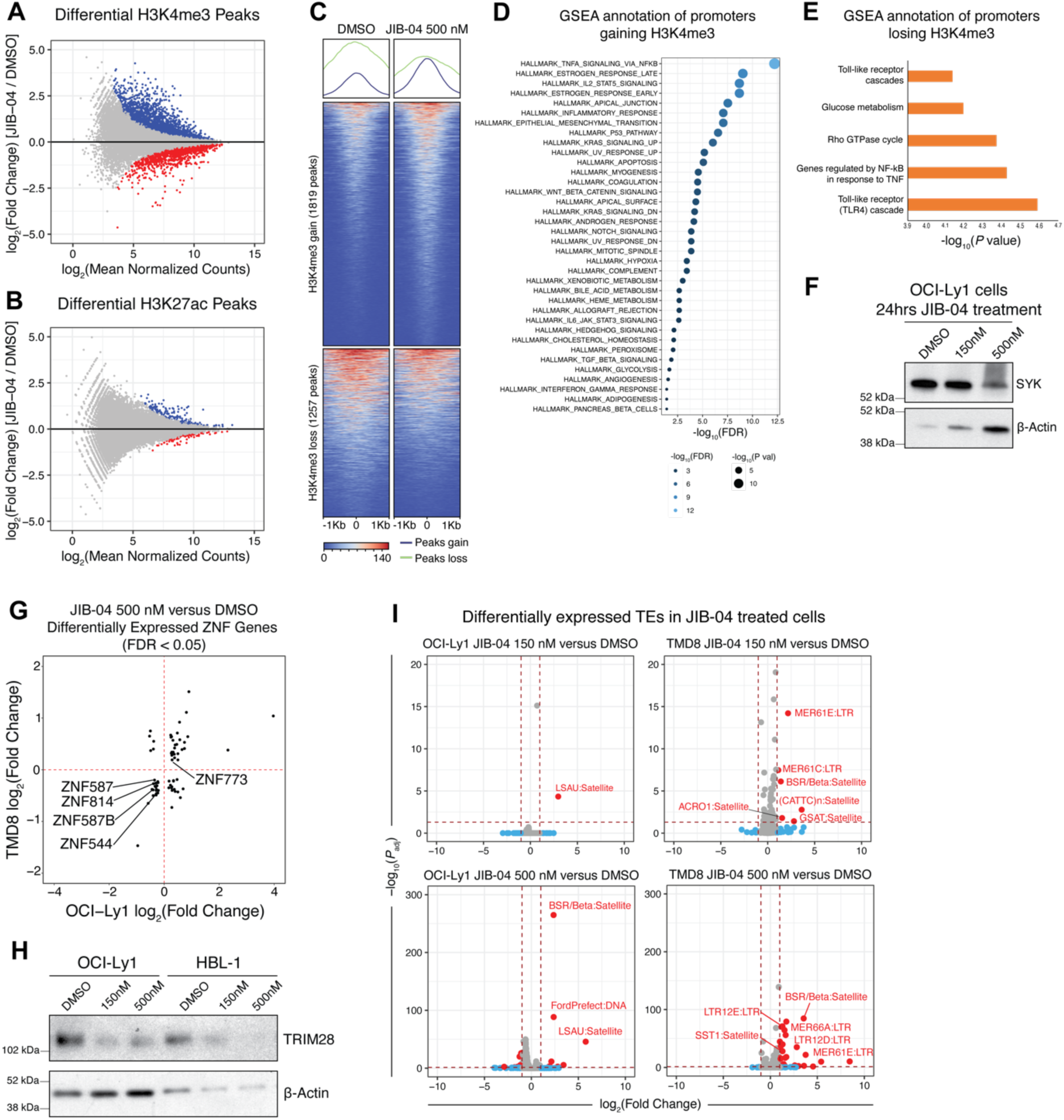
**A-B)** MA plots of ChIP-seq peaks exhibiting statistically significant differences for H3K4me3 **(A)** or H3K27ac **(B)** between JIB-04 (500 nM) and DMSO treated OCI-Ly1 cells. Peaks in JIB-04 treated cells with an FDR-adjusted *P* value < 0.05 and increased signal are in blue; decreased signal are in red. **C)** Heatmaps of H3K4me3 ChIP-seq signal from DMSO and JIB-04 treated OCI-Ly1 cells for all statistically significant peaks from **(A)**. Each row represents a ChIP-seq peak. Aggregate signal tracks across all peaks gaining or losing H3K4me3 are represented above the heatmap for each condition. **D-E)** Significant gene sets enriched within ChIP-seq peaks gaining **(D)** or losing **(E)** H3K4me3. Gene set enrichment analysis (GSEA) was performed on the set of gene promoters with a statistically significant change in H3K4me3 ChIP-seq signal. Gene sets were ranked by fold change in ChIP-seq signal between JIB-04 and DMSO treated conditions. **F)** Immunoblot for SYK in OCI-Ly1 cells after 24 hours of JIB-04 treatment. **G)** Fold change of significant (FDR-adjusted *P* value < 0.05) differentially expressed ZNF genes between 500 nM JIB-04 and DMSO treated OCI-Ly1 and TMD8 cells. Positive fold changes reflect up-regulation in JIB-04 treated cells. **H)** Immunoblot for TRIM28 in OCI-Ly1 and HBL-1 cells after 24 hours of JIB-04 treatment. **I)** Volcano plots of differentially expressed TEs between JIB-04 and DMSO treated OCI-Ly1 and TMD8 cells. Positive fold changes reflect up-regulation in JIB-04 treated cells relative to DMSO controls. TEs were considered statistically significant if FDR-adjusted *P* value < 0.05 and log_2_(fold change) > 0.75 or log_2_(fold change) < −0.75. Expression of TE families was quantified from RNA-seq data using the TEtranscripts software package^85^.

**Supplemental Figure 4.**
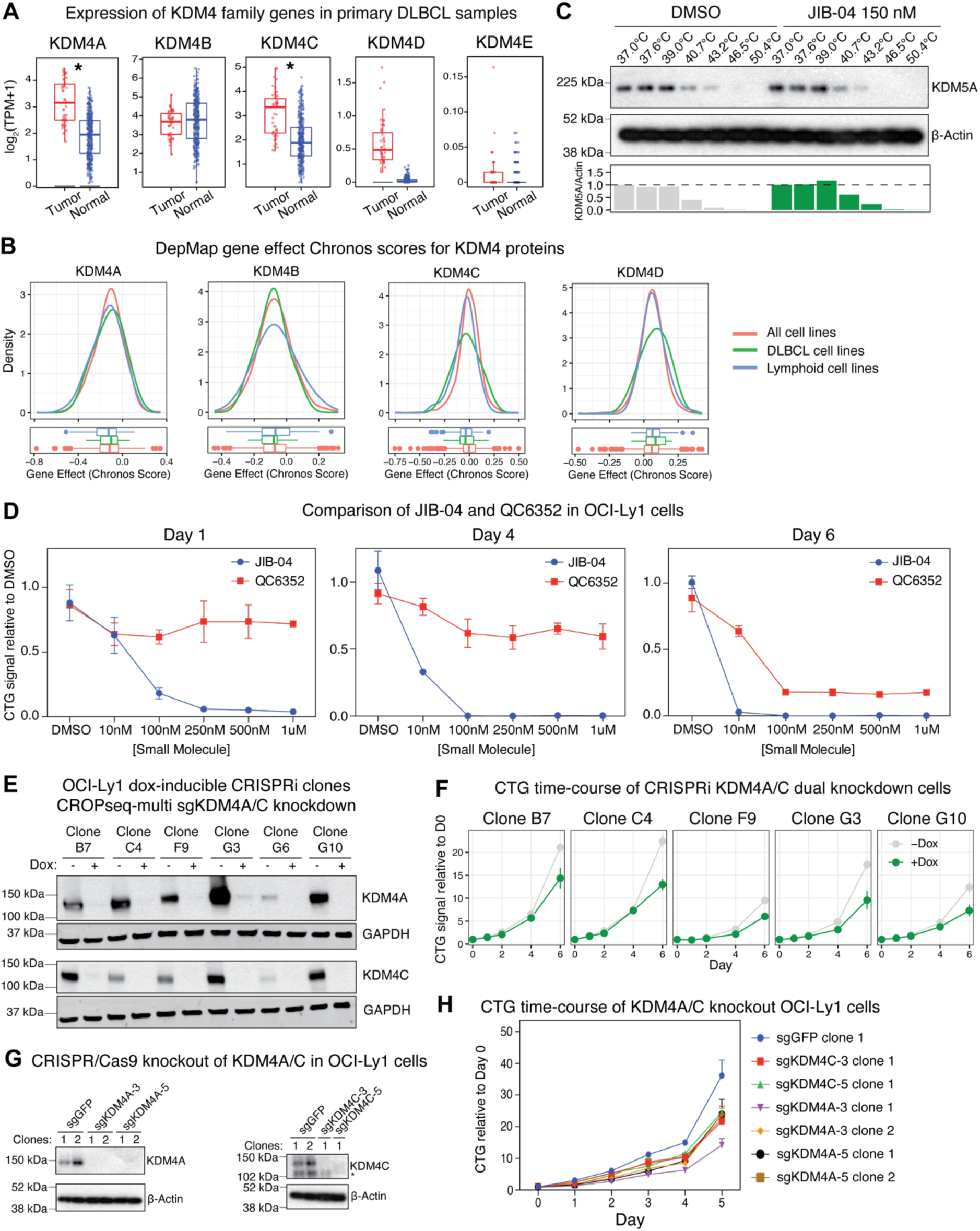
**A)** Expression of KDM4 genes in healthy and primary DLBCL patient samples obtained from the GEPIA2.0 database (PMID: 31114875). Each point reflects an individual patient. Asterisks represent FDR-adjusted *P* values < 0.01 (corrected for multiple hypothesis testing). **B)** Gene effect Chronos scores of KDM4 proteins from the 24Q4 release of DepMap. All cell lines n=1178, Lymphoid cell lines n=93, DLBCL cell lines n=16. **C)** Cellular thermal shift assay indicating lack of JIB-04 stabilization of KDM5A. Quantification of relative KDM5A protein signal (normalized to β-Actin loading control) is plotted for each sample below the blot. **D)** Dose titration of JIB-04 or QC6352 on OCI-Ly1 cells treated for one, four or six days. CellTiter-Glo luminescence values at each dose at each time point are normalized to DMSO controls. Error bars represent standard deviation over three biological replicates. **E)** Immunoblots of KDM4A and KDM4C in OCI-Ly1 cells after CRISPRi-mediated dual knockdown. A dox-inducible CRISPRi system was stably integrated into OCI-Ly1 cells by piggyBac transposition and multiple clones were derived. Each clone was infected with CROPseq-multi lentiviruses to express sgRNAs targeting the TSSs of KDM4A and KDM4C. Protein knockdown of KDM4A and KDM4C across all CRISPRi clones was assessed by immunoblotting after 2 weeks of 1 μg/mL dox treatment. **F)** CellTiter-Glo time-course of CRISPRi-mediated dual knockdown of KDM4A and KDM4C in various OCI-Ly1 clones from **(D)**. CellTiter-Glo luminescence values are normalized to the Day 0 baseline for each −/+dox condition. Error bars represent standard deviation across three biological replicates. **G)** Immunoblots of KDM4A (left panel) and KDM4C (right panel) proteins after CRISPR/Cas9-mediated knockout in OCI-Ly1 cells. Two different sgRNAs per gene were used to generate independent knockouts and an sgRNA targeting GFP was used as a non-targeting control. Asterisk represents a non-specific band. **H)** CellTiter-Glo time-course of KDM4A or KDM4C knockout OCI-Ly1 cells from **(F)**. CellTiter-Glo luminescence values are normalized to the Day 0 baseline for each sgRNA condition. Error bars represent standard deviation across three biological replicates.

**Supplemental Figure 5.**
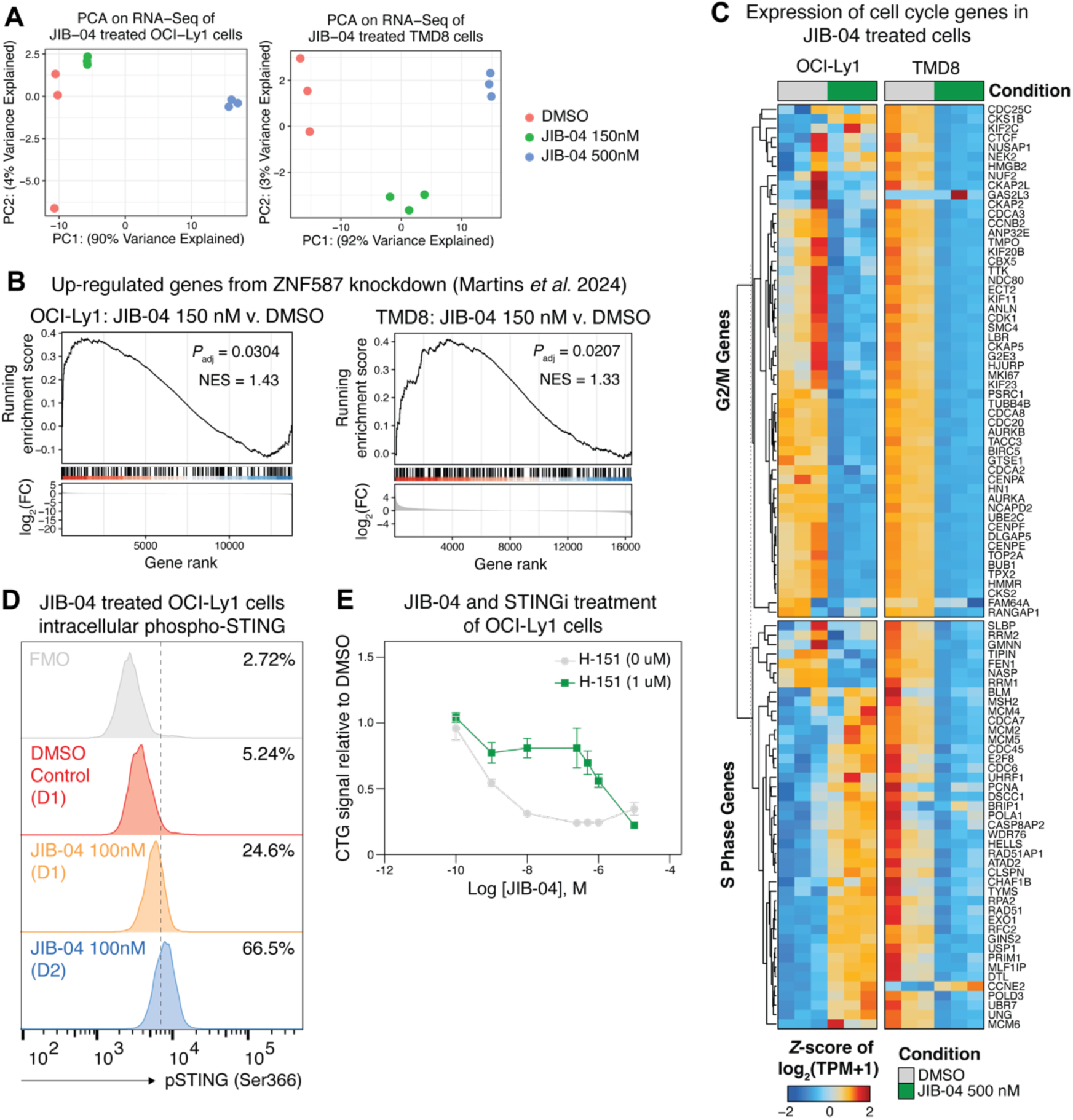
**A)** Principal component analysis (PCA) on RNA-seq data from DMSO and JIB-04 treated OCI-Ly1 (left) and TMD8 (right) cells. **B)** GSEA on 150 nM JIB-04 and DMSO treated OCI-Ly1 (top) and TMD8 (bottom) cells using a gene set comprising upregulated genes following knockdown of ZNF587 in DLBCL cell lines from Martins, *et al.* 2024 (PMID: 38345497). Genes were ranked by descending log_2_(fold change) from DESeq2. Positive NES scores reflect gene set enrichment within JIB-04 treated compared to DMSO treated cells. *P-*values were adjusted for multiple hypothesis testing. **C)** Expression of G2/M (top) and S (bottom) phase genes from Tirosh, *et al.* 2016 (PMID: 27124452) on DMSO and 500 nM JIB-04 treated OCI-Ly1 and TMD8 cells. Each column represents an independent biological replicate. Gene expression visualized in the heatmap are log_2_(TPM+1) data Z-score standardized across rows. TPM = transcripts per million. **D)** Intracellular flow cytometry time-course for pSTING (Serr366) on OCI-Ly1 cells treated with 100 nM JIB-04. **E)** Dose titration of JIB-04 +/− H-151 (STING inhibitor) after 48-hours treatment of OCI-Ly1 cells. CellTiter-Glo luminescence values are normalized to DMSO controls. Error bars represent standard deviation across three biological replicates.

**Supplemental Figure 6.**
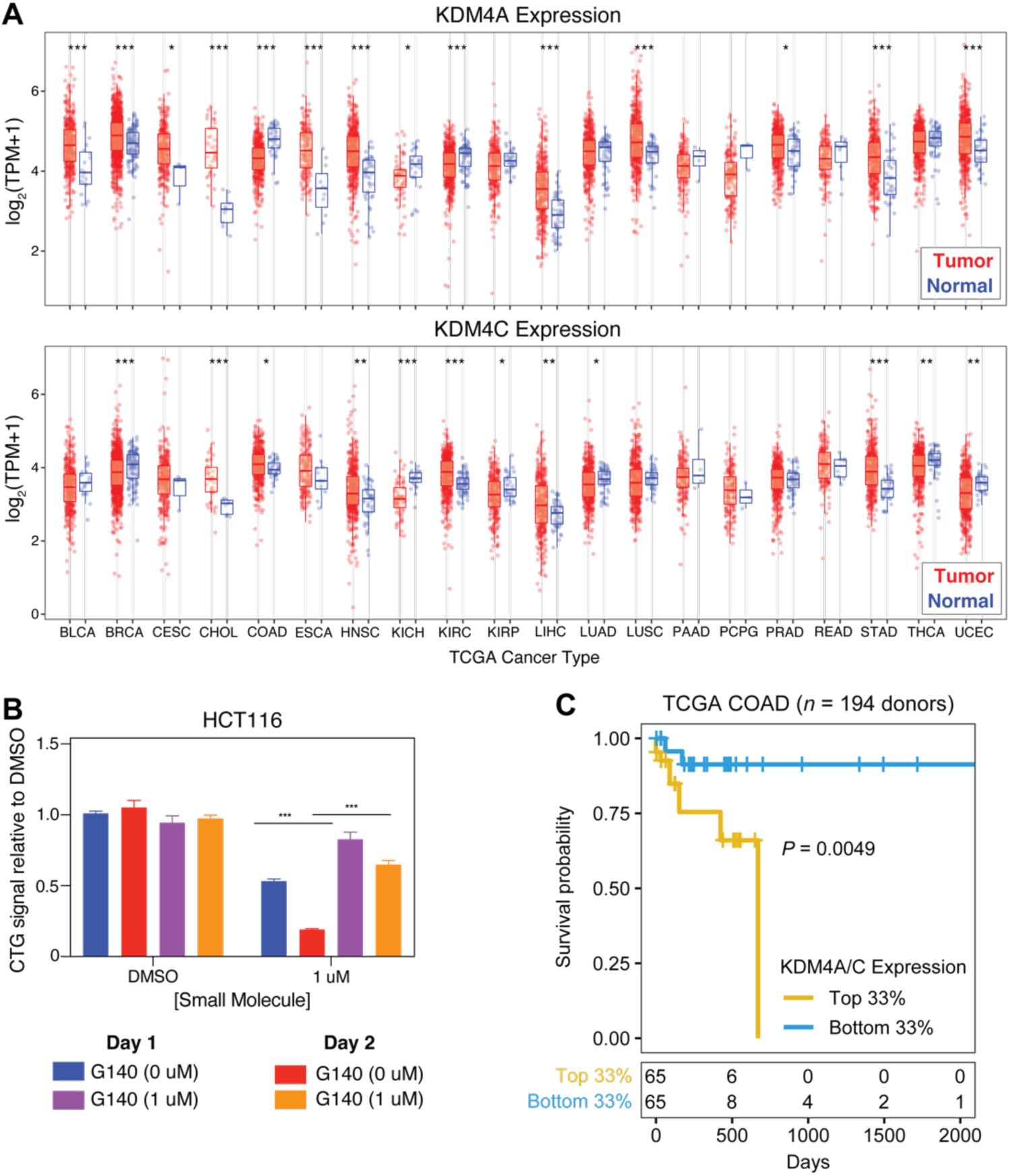
**A)** Expression of KDM4A (top) or KDM4C (bottom) in normal and tumor TCGA patient samples obtained from the TIMER2 database (PMID: 32442275)^86^. Each point represents an individual sample. Asterisks represent FDR-adjusted P values < 0.05 (corrected for multiple hypothesis testing) comparing tumor versus normal conditions for each TCGA cancer type via a Wilcoxon ranked sum *t*-test. **B)** HCT116 cells were treated with JIB-04 and G140 (cGAS inhibitor) for one or two days and CellTiter-Glo luminescence values were normalized to DMSO controls. Asterisks indicate FDR-adjusted *P* values < 0.05, comparing 0 versus 1 μM G140 treatment conditions via a Wilcoxon rank sum test. Error bars represent standard deviation across three biological replicates. **C)** Kaplan-Meier survival curves of TCGA COAD patients stratified by the top and bottom third of KDM4A/C expression.

**Supplemental Figure 7.**
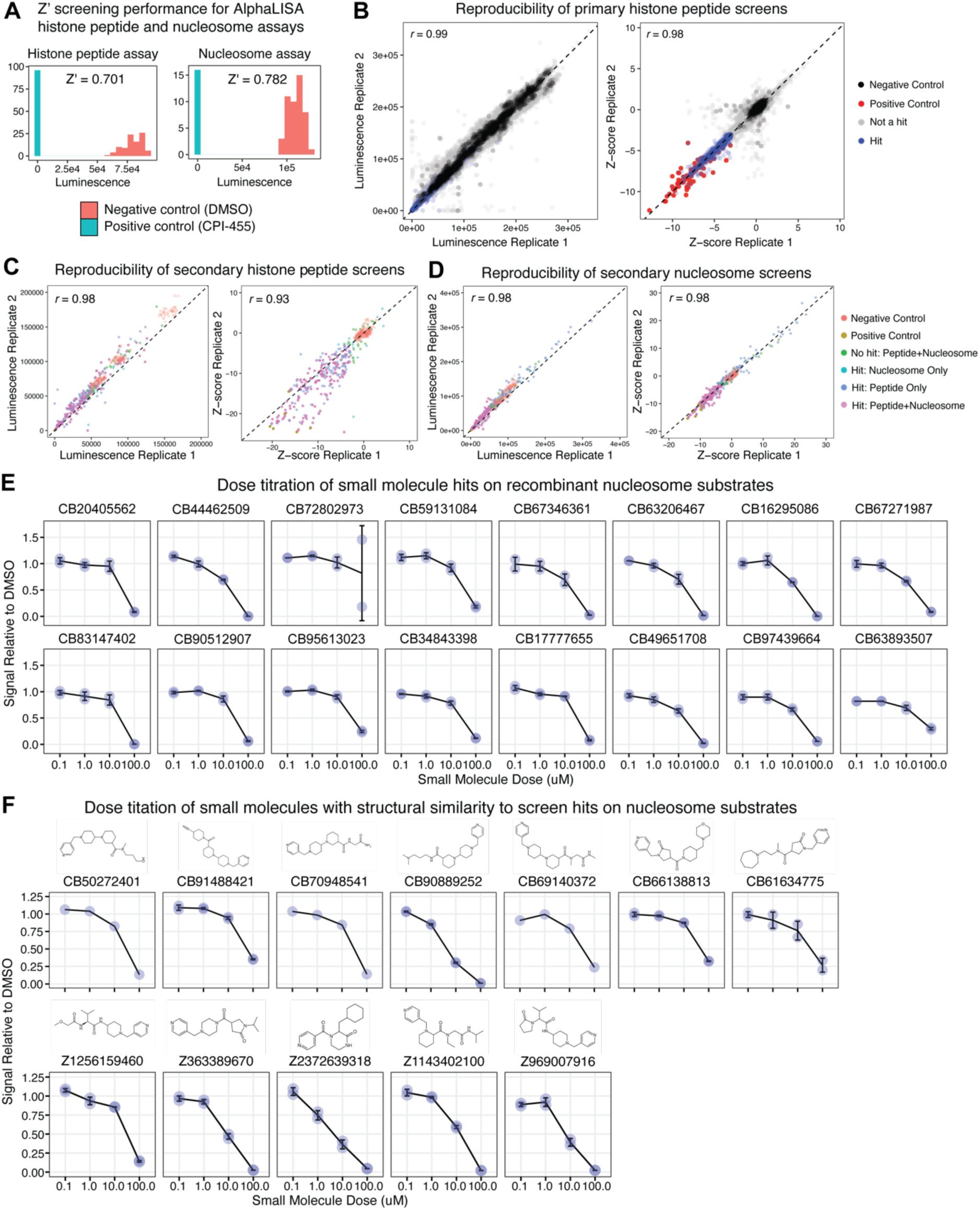
**A)** Distribution of luminescence values for negative (DMSO) and positive (CPI-455) controls in the AlphaLISA H3K9me3 histone peptide (left) and nucleosome (right) assay. Z-factor values for each assay are noted within each plot. **B)** Replicate reproducibility in the primary screen (from Figure 6B) of small molecule inhibitors for KDM4C using H3K9me3 peptide substrates. Raw luminescence values and Z-score normalized data for each replicate in the screen are respectively presented in the left and right plots. Z-scores for all small molecules and CPI-455 positive controls are calculated using the mean and standard deviation of the DMSO negative controls within each screening plate. Pearson correlations between screen replicates are noted within each plot. **C-D)** Replicate reproducibility in the secondary validation screen of small molecule inhibitors for KDM4C using H3K9me3 peptide **(C)** and nucleosome **(D)** substrates. Z-scores are calculated as described in **(B)** and Pearson correlations between each screen replicate are noted within each plot. **E-F)** Dose titration of small molecules hits in both histone peptide and nucleosome secondary validation screens **(E)**, and those that exhibit structural similarity to hits in Figure 6F. H3K9me3 demethylation activity was measured on nucleosome substrates. AlphaLISA luminescence values are normalized to DMSO controls. Error bars represent standard deviations across two replicates.

## SUPPLEMENTAL TABLES

**Table S1A:** Cell Titer-Glo data from OCI-Ly1 Cayman epigenetic small molecule screen

**Table S1B:** Cell Titer-Glo data from HBL-1 Cayman epigenetic small molecule screen

**Table S1C:** Processed Cayman epigenetic small molecule screen results

**Table S2A:** CROPseq-multi CRISPRi knockdown sgRNAs

**Table S2B:** CRISPR-Cas9 knockout sgRNAs

**Table S3:** Gene set of up-regulated genes upon knockdown of ZNF587 in DLBCL cells

**Table S4:** Processed KDM4i primary histone peptide screen data

**Table S5A:** Secondary screens of primary KDM4i small molecule hits

**Table S5B:** Processed KDM4i AlphaLISA TruHits screen data on prioritized primary screen hits

**Table S5C:** Top KDM4i small molecule hits

## SUPPLEMENTAL INFORMATION

### Resource Availability

#### Lead contact

Further information and requests for resources and reagents should be directed to and will be fulfilled by the lead contact, George Q. Daley.

#### Materials availability

All unique/stable reagents generated are available from the lead contact with a completed Materials Transfer Agreement.

#### Data and code availability

- Raw-omics data have been deposited at the NCBI Gene Expression Omnibus database and available from accession GSE286298
- All original code and processed data is available through the following Github repository: https://github.com/mnajia/Lymphoma_KDM4_Manuscript
- Any additional information required to reanalyze the data in this paper is available from the lead contact upon request

### Method Details

#### Animal studies

Immunocompromised NOD.Cg-*Prkdc^scid^ Il2rg^tm1Wjl^*/SzJ (NSG) mice (Jackson Laboratories cat. no. 005557) were housed at the Boston Children’s Hospital Animal Care facility following institutional guidelines. Male mice aged 6-8 weeks old received a subcutaneous injection of 1 million OCI-Ly1 cells (resuspended in 90% Matrigel and 10% PBS) to the left flank. Tumor size and overall body mass were measured three times a week. Mice were sacrificed once tumors reached 2.0 cm in diameter, at which point tumors and spleen were harvested. In separate experiments, mice were intravenously injected with 1 million OCI-Ly1 cells that constitutively expressed firefly luciferase. Inoculated animals were subjected to bioluminescence imaging (BLI) at regular intervals using the IVIS 200 system (PerkinElmer) following intraperitoneal injections of Vivoglo luciferin (Promega cat. no. P1043) at 150 mg/kg body weight. All animal experiments were performed under protocols approved by the Institutional Animal Care and Use Committee of Boston Children’s Hospital.

#### Cell culture

HBL-1, TMD8 and OCI-Ly1 cell lines were generously provided by Dr. Margaret Shipp and Dr. Catherine Wu of the Dana-Farber Cancer Institute. All other cell lines used in this study were obtained from ATCC or DSMZ. OCI-Ly1, OCI-Ly7, OCI-Ly3, OCI-Ly19 and HL-60 cells were cultured in IMDM + 10% FBS + Penicillin-Streptomycin. SUDHL-4, SUDHL-6, Karpas-422, TMD8, HBL-1, U-2932, Toledo, Pfeiffer, Farage, Kasumi-1 and U2940 cells were cultured in RPMI 1640 + 10% FBS + Penicillin-Streptomycin. Karpas-1106p cells were grown in RPMI 1640 + 20% FBS + Penicillin-Streptomycin. HEK293T and HCT116 cells were cultured in DMEM + GlutaMAX + 10% FBS. Cell lines were regularly monitored for mycoplasma through the Boston Children’s Hospital Stem Cell Core facility.

#### Cell-based phenotypic small molecule screen

Phenotypic screens in OCI-Ly1 and HBL-1 cells were performed using a library of 145 compounds that target various chromatin factors (Epigenetics Screening Library, Cayman Chemicals cat. no. 11076, Batch no. 0498084). Cells were cultured with compounds or DMSO vehicle controls for five days, and cell proliferation quantified with a CellTiter-Glo assay (Promega cat. no. G7572). In brief, a master mix of cells (200,000 cells / mL) was prepared for each line and 25 µL (5,000 cells) was added to each well of Nuncleon white polystyrene 96-well assay plates (Thermo Scientific cat. no. 136101) avoiding edge wells. Compounds were diluted in cell line specific media from a stock concentration of 10 mM to a working solution of 100 μM in 96-well plates, which were then serially diluted to 2 µM, 1 µM, and 200 nM. 25 µL of these solutions were administered to 25 µL cells to achieve the final assay concentrations of 1 µM, 500 nM, and 100 nM per compound. Each compound / dose in each cell line was assayed in triplicate. We also reserved 10 wells on each assay plate for 0.01% DMSO vehicle controls. In total, the screen amounted to 3,240 samples. Cell proliferation was quantified following five days of culture with CellTiter-Glo assays on a BioTek Synergy Neo microplate reader as per manufacturer’s protocol. The effect size of every compound / dose was determined by calculating the fold change of the mean CellTiter-Glo signal of compound-treated relative to DMSO-treated cells. Statistically significant compounds were identified using two-tailed Wilcoxon rank sum tests and a null distribution comprising all DMSO controls. *P* values were corrected for multiple hypothesis testing using the Benjamini-Hochberg method. Compounds were defined as hits from the screen if *P*-adjusted < 0.0001 and log_2_(fold change) < −1.

#### CellTiter-Glo viability time-course assay

We utilized CellTiter-Glo assays to determine the kinetics of cell viability upon chemical or genetic perturbations. First, 10,000 lymphoma cells or 5,000 HCT116 colorectal cells were seeded per well in Nuncleon white polystyrene 96-well assay plates (Thermo Scientific cat. no. 136101), avoiding edge wells with either DMSO, doxycycline, or small molecules at specific concentrations. Each condition was prepared in technical triplicate, and replicate plates were prepared at the start of the experiment for each timepoint in the time-course. The CellTiter-Glo assay (Promega cat. no. G7572) was performed according to the manufacturer’s protocol on a BioTek Synergy Neo microplate reader. The raw luminescence signal for each timepoint was normalized to the starting Day 0 baseline signal. Alternatively, data within a time-point was normalized to the DMSO controls for small molecule treatment experiments, or to the non-targeting sgRNA controls for CRISPR knockdown/knockout experiments. The following small molecules were utilized in this study: JIB-04 (Stem Cell Technologies cat. no. 73212), QC6352 (MedChem Express cat. no. HY-104048), EPZ011989 (Chemie Tek cat. no. CT-EPZ989), G140 (MedChem Express cat. no. HY-133916) and H-151 (MedChem Express cat. no. HY-112693).

#### Apoptosis analysis with Annexin V flow cytometry

Cells were treated with 0.1% DMSO or a dose titration of JIB-04 for 48 hours, after which the cells were harvested, washed with FACS buffer (PBS + 2% FBS) and stained with Annexin V with 7-AAD (BioLegend cat. no. 640930), according to the manufacturer’s protocol. Data was then collected using a Sony MA900 FACS and analyzed with FlowJo v10.10.

#### ChIP-seq

For ChIP-Seq experiments, 10 million OCI-Ly1 cells were fixed with 1% formaldehyde for 10 minutes at room temperature before quenching with 0.125 M glycine. Cells were then lysed in ChIP buffer (0.6% SDS, 10 mM EDTA, and 50 mM Tris-HCl, pH 8.1) and cross-linked chromatin was sonicated to obtain DNA fragments of 300-800 bp. Samples were centrifuged (10 min, 4°C, 13,000 rpm) to clarify, and the chromatin supernatant was diluted 6-fold in ChIP dilution buffer (0.01% SDS, 1.1% Triton X-100, 1.2 mM EDTA, 16.7 mM Tris-HCl pH 8, and 167 mM NaCl) with protease inhibitors (Roche Diagnostics). Diluted chromatin was incubated with 5 μg of primary antibody overnight at 4°C with agitation. Antibodies were captured using 50 μL of Dynabeads (ThermoFisher Scientific). The beads were sequentially washed once on a magnet with ChIP wash buffer 1 (0.1% SDS, 1% Triton X-100, 2 mM EDTA, 20 mM Tris-HCl pH 8, and 150 mM NaCl), then ChIP wash buffer 2 (0.1% SDS, 1% Triton X-100, 2 mM EDTA, 20 mM Tris-HCl pH 8, and 500 mM NaCl), then ChIP wash buffer 3 (0.25 M LiCl, 1% IGEPAL-CA630, 1% deoxycholic acid, 1 mM EDTA, 10 mM Tris pH 8), and finally then ChIP wash buffer 4 (10 mM Tris-HCl, 1 mM EDTA). Chromatin was eluted using ChIP Elution Buffer (1% SDS, 100mM NaHCO3). Crosslinks were reversed by incubation at 65°C overnight, and proteins were then digested with proteinase K. DNA was recovered by phenol-chloroform extraction and ethanol precipitation in presence of 20 µg of glycogen (Sigma Aldrich). ChIP-seq libraries were prepared using NEBNext DNA Library Prep Master Mix Set for Illumina (NEB cat. no. E6040). Libraries were single-end indexed and sequenced on an Illumina HiSeq 2000 to generate 50 bp single end reads. Histone antibodies for ChIP-Seq were chosen based on specificity data from http://www.histoneantibodies.com/ and the antibodies used in this study include, H3K9me2 (Abcam cat. no. ab1220, RRID:AB_449854), H3K9me3 (Active Motif cat. no. 39161, RRID:AB_2532132 and Abcam cat. no. ab8898, RRID:AB_306848), H3K4me3 (EpiGentek cat. no. A-4033-050, RRID:AB_2920607), and H3K27Ac (Active Motif cat. no. 39135, RRID:AB_2614979).

#### ChIP-seq analysis

ChIP-seq libraries were processed using the nf-core/chipseq pipeline v2.0.0 (https://doi.org/10.5281/zenodo.3240506) implemented in the NextFlow software package (v23.10.1). In brief, single-end reads were trimmed of adaptor sequences with trim-galore (v0.6.7) and aligned with BWA (v0.7.17-r1188) to the GRCh38 assembly of the human genome. Reads aligning to the ENCODE exclusion list (<u>ENCFF356LFX</u>) were omitted. Duplicate reads were removed with Picard (v2.27.4) with the MarkDuplicates tool. Peaks for H3K4me3 and H3K27ac ChIP-seq experiments were identified with MACS2 (v2.2.7.1) using the BAM files of the input ChIP-seq control and histone ChIP-seq samples with the following command line parameters: -- keep-dup all --broad --broad-cutoff 0.1 --gsize 2701262066. We created bigwig files for visualization using bamCoverage from the deepTools package (v3.5.1) with the following command line options: --binSize 20 --normalizeUsing RPKM --smoothLength 60 --extendReads 150 --centerReads.

To identify differential H3K4me3 and H3K27ac ChIP-seq peaks, we created consensus peaksets for each ChIP-seq experiment across all samples and counted aligned reads within the consensus peakset using featureCounts from the subread package (v2.0.1). We then utilized these count matrices to determine statistically significant peaks varying between DMSO and JIB-04 treated conditions using DESeq2 (v1.40.2) with an FDR-based *P* value adjustment method and alpha = 0.05. Differential peaks were considered statistically significant if *P* adjusted < 0.05.

We used chromVAR to infer TF activity in the consensus set of H3K4me3 or H3K27ac ChIP-seq peaks. First, we utilized a set of non-redundant TF “motif archetypes” (https://resources.altius.org/~jvierstra/projects/motif-clustering-v2.1beta/)^87^, which represent clustered motifs that have been deduplicated based on similarity across all known human TF motifs. The motif archetypes were imported into R as PWMatrices. We then determined motif matches within the consensus set of H3K4me3 or H3K27ac ChIP-seq peaks using the “matchMotifs” function from the motifmatchr package in R. We computed GC biased-corrected deviations via the chromVAR “deviations” function using the raw counts matrix across consensus peaks and motif matches as input. We identified motif archetypes that exhibited statistically significant differences between DMSO and JIB-04 treated conditions using the “computeVariability” chromVAR function and an FDR-adjusted *P* value cut off of 0.05. We then plotted the chromVAR deviation z-scores for all significant motif archetypes in H3K4me3 and H3K27ac peaks across all samples using the ComplexHeatmap package in R.

To identify genomic regions with differential H3K9me2 or H3K9me3 enrichment in JIB-04 treated cells, we performed window-based analyses using the csaw package (v1.34.0)^88^ in R due to the broad distribution of these histone marks on chromatin. Aligned reads were extended to 200 bp and counted into 2kb windows for each ChIP-seq library. Reads were also counted into 10kb bins, and the median average abundance of all bins was used as a global estimate of the background abundance. Windows were filtered to retain only those with a two-fold or greater increase in the average abundance above the scaled background estimate. Counts from the remaining windows were tested for significant ChIP signal enrichment using edgeR (v3.42.4). Briefly, an abundance-dependent trend in the NB dispersions was fitted to all windows, using the estimateDisp function. A GLM was fitted to the counts for each window using the trended NB dispersion. The quasi-likelihood (QL) dispersion was estimated from the GLM deviance. An abundance-dependent trend was robustly fitted to the QL dispersions across all windows, and the QL dispersion estimate for each window was shrunk to this trend. Finally, a *P* value for differential ChIP signal in each window was computed using the QL F-test.

#### RNA Sequencing

Total RNA from JIB-04 treated OCI-Ly1 and TMD8 cells were isolated using the RNeasy Plus Micro kit (Qiagen cat. no. 74034) per the manufacturer’s protocol. Purified RNA was subjected to ribosomal depletion, using the NEBNext rRNA Depletion Kit (New England Biolabs cat. no. E7770L). RNA-Seq libraries were prepared in technical triplicate using the NEBNext Ultra RNA II Library Prep Kit (New England Biolabs cat. no. E6310L) and paired-end sequenced with 101 cycles each read on an Illumina HiSeq.

#### Analysis of gene expression from RNA-seq data

Paired-end RNA-seq reads were pseudoaligned to the hg19 reference transcriptome with Kallisto (version 0.46.0)^89^ to quantify transcriptome-wide gene expression. Transcript-level estimates were summed using the tximport package (version 1.28.0) in R (version 4.3.1). Differential expression was performed using the DESeq2 package (version 1.40.2)^90^ on the estimated counts matrix from Kallisto. Statistically significant genes varying between two conditions were identified using an FDR-based *P* value adjustment method and alpha = 0.05. We defined genes as significantly differentially expressed if *P* adjusted < 0.05 and log_2_(fold-change) > 1 or log_2_(fold-change) < −1. Gene set enrichment analysis (GSEA) using Hallmark gene sets was performed using the clusterProfiler package (version 4.8.2) and an unfiltered gene list from DESeq2 ranked by descending log_2_(fold-change) as input. Gene sets were considered significantly enriched if an FDR-adjusted *P* value < 0.05. Furthermore, we downloaded RNA-seq data from Martins, *et al.* 2024^64^ (GSE229468), processed it with Kallisto and performed differential gene expression analysis with DESeq2, as described above. We created a gene set that comprised the set of conserved genes that were up-regulated (log_2_(fold change) > 0.75 and FDR-adjusted *P* value < 0.05) between shZNF587 and shControl conditions in OCI-Ly7 and U-2932 cells. We then utilized this gene set to perform GSEA on our RNA-seq data comparing JIB-04 versus DMSO treated OCI-Ly1 and TMD8 cells.

#### Analysis of transposable element expression from RNA-seq data

RNA-seq reads were mapped to the hg38 genome using STAR aligner^91^ with command line flags “--winAnchorMultimapNmax 200 --outFilterMultimapNmax 200” and the resulting BAM files were used with TEtranscripts^85^ to quantify TE expression. Curated GTF files of TE annotations in the hg38 assembly of the human genome were downloaded from the Hammell Lab website for TEtranscript quantification. Differential expression was subsequently performed with DESeq2 (v1.10.1) comparing JIB-04 and DMSO treated cells. Differentially expressed TEs were defined as log_2_(fold change) > 0.75 or log_2_(fold change) < −0.75 and FDR-adjusted *P* value < 0.05. We plotted H3K9me3 ChIP-seq signal over differentially expressed satellite regions () using deepTools (v3.5.1) and ChIP-seq bigwig files as input.

#### Assaying inhibition of KDM4 activity on H3K9me3 nucleosome substrates with AlphaLISA

KDM4A/C enzymatic activity / inhibitors thereof were assayed by the relative conversion of nucleosomal H3K9me3 substrate to H3K9me2 product.

KDM4A: In brief, 2.5 uL of each compound titration was prepared in KDM4A reaction buffer (10 mM Tris-HCl pH 7.5 + 0.01% Tween-20 + 0.01% BSA) supplemented with 4% DMSO and combined in a 384-well plate with 5 uL of 8 nM KDM4A (Uniprot O75164; aa1-350, N-terminally GST tagged), and pre-incubated for 15 minutes at 23°C. A 2.5 uL mix of 400 mM Ascorbic acid, 50 mM Alpha ketoglutarate, 45 mM Ammonium Iron, and 40 nM H3K9me3 biotinylated nucleosome (5’ DNA biotinylated; EpiCypher cat. no. 16-0315) in the KDM4A reaction buffer was added to each well, and reactions incubated for 30 minutes at 23°C. The enzyme reaction product was detected using 10 uL of anti-H3K9me2 (RevMab cat. no. 31-1059-00, RRID:AB_2716382) prepared in KDM4A quench buffer (10 mM Tris pH 7.5 + 0.01% Tween-20 + 0.01% BSA + 625 mM NaCl + 20 mM EDTA). Following a 30-minute incubation, a 5 uL mixture of 30 mg/mL Protein A AlphaLISA Acceptor beads (PerkinElmer cat. no. AL101) and 100 mg/mL Streptavidin Donor beads (PerkinElmer cat. no. 6760002) prepared in KDM4A bead buffer (10 mM Tris pH 7.5 + 0.01% Tween-20 + 0.01% BSA) was added. The plate was incubated for 60 minutes at 23°C in subdued lighting, and AlphaLISA signal was measured on a PerkinElmer 2104 EnVision (680 nm laser excitation, 570 nm emission filter +/− 50 nm bandwidth).

KDM4C: In brief, 5 uL of each compound titration was prepared in KDM4C reaction buffer (10 mM Tris-HCl pH 7.5 + 0.01% Tween-20 + 0.001% Casein + 1 mM TCEP) supplemented with 3% DMSO and combined in a 384-well plate with 5 uL of 15 nM KDM4C (Uniprot Q9H3R0; aa1-350, N-terminally GST tagged) and pre-incubated for 15 minutes at 23°C. A 5 uL mix of 300 mM Ascorbic acid, 75 mM Alpha ketoglutarate, 90 mM Ammonium Iron, and 15 nM H3K9me3 biotinylated nucleosome (5’ DNA biotinylated; EpiCypher cat. no. 16-0315) in the KDM4C reaction buffer was added to each well, and the reactions were incubated for 30 minutes at 23°C. The enzyme reaction product was detected using 5 uL anti-H3K9me2 (RevMab cat. no. 31-1059-00, RRID:AB_2716382) prepared in KDM4C quench buffer (10 mM Tris pH 7.5 + 0.01% Tween-20 + 0.001% Casein + 625 mM NaCl + 20 mM EDTA). Following a 30-minute incubation, a 5 uL mixture of 12.5 mg/mL Protein A AlphaLISA Acceptor beads (PerkinElmer cat. no. AL101) and 40 mg/mL Streptavidin Donor beads (PerkinElmer cat. no. 6760002) prepared in KDM4C bead buffer (10 mM Tris pH 7.5 + 0.01% Tween-20 + 0.001% Casein) was added. The plate was incubated for 60 minutes at 23°C in subdued lighting, and AlphaLISA signal was measured as above.

#### Cellular thermal shift assay

Protein targets of JIB-04 were identified with cellular thermal shift assays as previously described^92^. Briefly, OCI-Ly1 cells were treated with 150 nM JIB-04 (Stem Cell Technologies cat. no. 73212) or DMSO for 1 and 3 hours. Drug-treated cells were heated (37-53.6°C) for 3 minutes on a thermal cycler then immediately snap-frozen in liquid nitrogen. Cells were lysed with two freeze-thaw cycles using liquid nitrogen and a thermal cycler set to 25°C. The soluble protein fraction was isolated by centrifuging the cellular lysate at 20,000g for 20 min at 4°C. Immunoblotting was performed for KDM4A, KDM4C, KDM5A and β-Actin.

#### Whole cell protein extraction and immunoblotting

To extract protein for immunoblotting, cells were resuspended in RIPA buffer supplemented with a protease inhibitor cocktail (Sigma cat. no. 4693159001) and incubated on ice for 15 minutes followed by sonication (30 seconds on, 30 seconds off for 15 cycles). The sonicated samples were centrifuged at 20,000g for 20 minutes at 4°C and the supernatant was collected as the whole cell extract. Protein concentrations were quantified with a BCA assay (Thermo Scientific cat. no. 23225), according to the manufacturer’s protocol. Cellular protein lysate was denatured in Laemmli Sample Buffer (BioRad cat. no. 1610737) for 10 minutes at 90°C then 20 μg of protein was run on an SDS-PAGE gel (BioRad cat. no. 4561023) for 50 minutes at 180 V. Protein was then transferred to a PVDF membrane using a wet transfer method for 3 hours at a constant current of 50 mA. The PVDF blots were then blocked with 5% milk (BioRad cat. no. 1706404) in TBS-T with gentle agitation for 1 hour at room temperature, and then incubated with primary antibodies overnight at 4°C. Primary antibodies used in this study include KDM4A (Abcam cat. no. ab191433, 1:1000 dilution), KDM4C (R&D Systems cat. no. AF6430, RRID:AB_10718990, 1:1000 dilution), KDM5A (Abcam cat. no. ab194286, RRID:AB_2889152, 1:1000 dilution), TRIM28 (Abcam cat. no. ab22553, RRID:AB_447151, 1:500 dilution), IKZF1 (Cell Signaling Technology cat. no. 5443S, RRID:AB_10691693, 1:1000 dilution), IKZF3 (Novus BIologicals cat. no. NBP2-24495, RRID:AB_3096994, 1:1000 dilution), SYK (Cell Signaling Technology 2712S, RRID:AB_2197223, 1:1000 dilution), TBP (Cell Signaling Technology 8515S, RRID:AB_10949159, 1:10,000), Apoptosis Western-Blot Cocktail (Abcam cat. no. ab136812, 1:500), GAPDH (Santa Cruz Biotech cat. no. SC-32233, RRID:AB_627679, 1:5000 dilution), β-Actin (Cell Signaling Technology cat. no. 3700S, RRID:AB_2242334, 1:10000 dilution), and Histone 3 (Active Motif 39451, RRID:AB_2793242, 1:5000 dilution). The blots were then stained with a Donkey anti-mouse IRDye 680RD secondary antibody (Li-Cor cat. no. 926-68072, RRID:AB_10953628, 1:10000 dilution) and imaged on a Licor fluorescence imager, or stained with an HRP-conjugated secondary antibody (1:5000 dilution) and visualized with HRP substrate SuperSignal West Dura (Thermo Scientific cat. no. 34075). Immunoblots were analyzed in Fiji v2.14.0/1.54f.

#### ZNF587 co-immunoprecipitation

We determined if ZNF587 interacts with KDM4 proteins in DLBCL cells based on established co-immunoprecipitation protocols^93^, but with some modifications. First, 10 million OCI-Ly1 cells were lysed on ice for 30 minutes in a buffer consisting of 10 mM HEPES, 10 mM KCl, and 0.05% NP-40 supplemented with a protease inhibitor cocktail (Sigma cat. no. 4693159001). The lysate was centrifuged for 15 minutes at 4°C at 16,000g and then the protein concentration was quantified with a BCA Protein Assay kit (Fisher Scientific cat. no. 23225). Dynabeads (Fisher Scientific cat. no. 14311D) were coupled to an anti-ZNF587 antibody (Thermo Scientific cat. no. MA5-22800, RRID:AB_2605893) or an IgG control antibody (BioLegend cat. no. 400101, RRID:AB_2891079) for 18 hours at 37°C on an orbital rotator. The antibody-coupled beads were incubated with 2 mg of total protein lysate for 24 hours at 4°C on an orbital rotator. The beads were then washed as previously described^93^ and the protein immunoprecipitate was eluted from the beads with SDS and incubation at 98°C. The protein immunoprecipitate from IgG and anti-ZNF587 pull-downs was assayed via immunoblots for ZNF587 (Thermo Scientific cat. no. MA5-22800, RRID:AB_2605893, 1:1000 dilution), KDM4A (Abcam cat. no. ab191433, 1:1000 dilution), KDM4C (R&D Systems cat. no. AF6430, RRID:AB_10718990, 1:1000 dilution), and GAPDH (Santa Cruz Biotech cat. no. SC-32233, RRID:AB_627679, 1:5000 dilution), as previously described.

#### B cell clonogenic potential assay

Lymphoma cells were resuspended in MethoCult H4531 (StemCell Technologies cat. no. 04531) supplemented with DMSO or small molecules. Cells were then cultured in 3.5 cm dishes in a humidified chamber inside a 37°C incubator for 14 days (n=4 replicate assays per condition), at which point colonies were scored by an investigator blinded to the treatment conditions.

#### Molecular cloning

All plasmids utilized in this study were designed in Benchling. Lentiviral vectors for multiplexed sgRNA expression were generated using the CROPseq-multi system^94^. First, we empirically tested the activity of individual CRISPRi sgRNAs pulled from the Broad Institute’s Dolcetto Library^95^ and verified protein knockdown by immunoblotting. Highly active CRISPRi sgRNAs were then prioritized for multiplexed gene knockdown experiments. Dual sgRNA inserts [BsmBI-<spacer #1>-<sgRNA scaffold>-<Gln tRNA>-<spacer #2>-BsmBI] were purchased as gene fragments from Twist Biosciences. The sgRNA inserts were cloned into CROPseq-multi-Puro (Addgene #216217) via Golden Gate Assembly at 1:100 (vector:insert) molar ratio using Esp3I (Fisher Scientific cat. no. ER0451) and T7 ligase (Enzymatics cat. no. L6020L) with the following thermal cycling protocol: 15 cycles of (37°C for 5 min, 20°C for 5 min). All sgRNA sequences used in this study are documented in **Supplemental Table S2.**

All assembled plasmids were purified with a DNA Clean & Concentrator column (Zymo cat. no. D4033) prior to transformation into NEB Stable Competent *E. coli* (cat. no. C3040H). Transformed *E. coli* were grown overnight at 30°C on agar plates with 50 µg/mL carbenicillin selection. Individual colonies were picked for liquid culture in LB media supplemented with 100 µg/mL ampicillin and plasmid DNA was subsequently isolated using a Qiagen Miniprep Kit (Qiagen cat. no. 27106). Plasmid sequences were fully verified by whole plasmid sequencing (Primordium Labs, Inc). All plasmids have been deposited with Addgene.

#### Construction of inducible CRISPR-interference cell lines

OCI-Ly1 cells were engineered to express dox-inducible CRISPRi machinery through piggyBac transposition. First, 1 million cells were nucleofected with 1 µg of piggyBac transposase plasmid (SBI cat. no. PB200A-1) and 5 µg of PB-NDi-ZIM3-KRAB-dCas9 transposon plasmid using SF cell line buffer (Lonza cat. no. V4XC-2012) and pulse code FF-100 on a Lonza 4D-Nucleofector. Cells were allowed to recover and expand for five days. Doxycycline was subsequently administered at a final concentration of 1 µg/mL for 48 hours to induce expression of the ZIM3-KRAB-dCas9-P2A-mCherry transgene. Single cells with high mCherry expression were isolated using a Sony MA900 FACS into 96-well plates to derive clones. We prioritized clones with no transcriptional leakiness (evidenced by flow cytometry for mCherry in the absence of dox treatment) and for uniform induction of the ZIM3-KRAB-dCas9-P2A-mCherry transgene by flow cytometry 24 hours after administration of 1 µg/mL dox. Prioritized clones were then expanded, banked and used for subsequent knockdown experiments.

#### Lentivirus production and infection of sgRNA vectors

Lentivirus was produced for each CRISPR and sgRNA expression vector by transfection of pMD2.G (Addgene #12259), psPAX2 (Addgene #12260), and the transfer plasmid (2:3:4 ratio by mass, 3 µg total) into HEK293FT cells using Lipofectamine 3000 (Thermo Scientific cat. no. L3000015). Viral supernatants were harvested 48 hours after plasmid transfection, centrifuged at 1000g for 5 minutes, and then filtered through 0.45 μm PVDF filters (Millipore cat. no. SLHVR04NL). Inducible CRISPRi cells were seeded into 6-well plates at 250,000 cells/well and infected with lentiviruses at MOI=0.25 in the presence of 8 μg/mL polybrene (Milipore Sigma cat. no. TR-1003-G). After 48 hours of incubation with lentiviruses, the cells were selected with 1 μg/mL puromycin (Life Technologies cat. no. A1113803) until the no-infection control cells completely died (∼5 days). Fully selected cells were expanded, banked and then used for downstream experiments.

#### Generation and validation of CRISPR knockout cell lines

We generated CRISPR knockouts via lentiviral delivery of sgRNAs into cells stably expressing spCas9 (for KDM4A and KDM4C knockouts). We generated independent knockouts for each gene with distinct sgRNAs and the sgRNA sequences used to knockout each gene are documented in **Supplemental Table S2**. We FACS-isolated clones from all edited cell lines and validated the knockouts by immunoblotting, as described in the “Whole cell protein extraction and immunoblotting” section.

#### Intracellular staining of pSTING and pH2AX foci

OCI-Ly1 and HBL-1 cells treated with 0.02% DMSO or 100 nM QC6352 for 48 hours were fixed in 4% PFA for 15 minutes at room temperature on a shaker set to 400 RPM. The cells were then washed twice with FACS buffer (PBS + 2% FBS) and kept at 4°C until ready for permeabilization. Cells were permeabilized on ice for 15 minutes in 100% methanol followed by two washes with FACS buffer. We then stained the permeabilized cells with DAPI and primary conjugated antibodies targeting phospho-STING (Ser366) (Cell Signaling Technology cat. no. 41622S, RRID:AB_2799204) or phospho-H2A.X (Ser139) (Cell Signaling Technology cat. no. 9720S, RRID:AB_10692910) at 1:50 dilution in FACS buffer for 60 minutes at room temperature sheltered from light. The stained cells were then washed with FACS buffer prior to analysis. Cells stained for intracellular pSTING were analyzed on a CytoFLEX S flow cytometer (Beckman Coulter) with factor-minus-one staining controls and the flow cytometry data was analyzed in FlowJo. Cell stained for pH2A.X were imaged in a Cellvis 384-well glass-bottom plate using a Yokogawa CellVoyager 7000 confocal imager (40x, 0.5µm Z steps, 405/640nm excitation, 445/676nm emission, 1×1 binning, 16 fields per well). FIJI/ImageJ was used to identify the Z level with the maximal DAPI signal. Nuclei were identified by applying the Versatile (fluorescent nuclei) trained model of the Stardist algorithm^96^ to flatfield-corrected DAPI images at the selected Z level followed by removal of overlapping or edge-touching ROIs. pH2A.X speckles were identified as local maxima (prominence ≥ 150) and the distributions of pH2A.X speckle intensities across treatment conditions were plotted in R (version 4.2.2).

#### Cell cycle analysis via flow cytometry of EdU incorporation

OCI-Ly1 and HBL-1 cells were treated with 0.02% DMSO or 100 nM QC6352 for 48 hours (n=3 biological replicates per condition). The cells were then pulse labeled with 10 μM Edu for 1 hour prior to fixation. EdU incorporation was detected using an EdU Click Proliferation Kit (BD Biosciences cat. no. 565456, RRID: AB_2869678) according to the manufacturer’s instructions. Cells were resuspended in FACS buffer (PBS + 2% FBS), stained with 2 μg/mL DAPI, and subsequently analyzed on a Sony MA900 FACS. Flow cytometry data was analyzed using FlowJo v10.10.

#### Survival analysis

For survival analysis, we downloaded processed RNA-Seq data of colorectal cancer patients^97^ from the GDC portal (https://gdc.cancer.gov/about-data/publications/coadread_2012) and patient survival data from the RTCGA.clinical Bioconductor package (v20151101.28.0) in R. We matched patients between both datasets using TCGA IDs. We then computed row-wise expression *z*-scores for *KDM4A* and *KDM4C* genes. Next, we took the column means of this matrix to get an average *z*-score across KDM4 genes and then identified the top 33% and bottom 33% of patients based on this mean standardized expression. We computed Kaplan-Meier curves using the kmTCGA function from the RTCGA package (v1.28.0) in R.

#### High-throughput small molecule screens to identify novel KDM4 inhibitors

To identify novel chemical matter targeting KDM4 demethylases, we performed small molecule screens using the ChemBridge 2020 library from the ICCB-Longwood Screening facility at Harvard Medical School (https://iccb.med.harvard.edu/chembridge-2020). The library consists of 50,000 compounds at a stock concentration of 10 mM in DMSO arranged across 143 384-well plates. DMSO vehicle controls were present on every plate in column 23 and column 24 was empty. To perform the primary screen, we screened 10 library plates per day in technical duplicate using a recombinant histone peptide AlphaLISA assay. In brief, columns 1-24 of 384-well assay plates (PerkinElmer cat. no. 6005359) were prefilled with 10 uL of 15.8 nM recombinant human KDM4C protein and spun down. Next, 100 nL of each compound was echo-transferred from the compound library plate to each assay plate. Then, 1 uL of 9 mM positive control compound, CPI-455 was added to each well in column 24 using a Multidrop Combi Reagent Dispenser (ThermoFisher Scientific). The plates were incubated for 30 minutes at room temperature. Next, 2.5 uL of a cofactor and peptide substrate solution was added such that the concentrations were as follows in 12.5 uL: 100 uM Ascorbic Acid, 5 uM Alpha ketoglutarate, 50 uM Iron, and 20 nM H3K9me3 peptide. The plates were then spun down and incubated for 30 minutes at room temperature. Subsequently, 2.5 uL of anti-H3K9me1 antibody was added to each well. Once again, the plates were spun down and incubated for 30 minutes at room temperature. Lastly, 15 uL of an AlphaLISA acceptor and donor bead solution was added to each well. The final assay volume per well was 30 uL. Assay plates were covered with TopSeal-A Black seals (PerkinElmer cat. no. 6050173), spun down, and incubated sheltered from light for 45 minutes prior to reading luminescence on a PerkinElmer Envision instrument in Alpha mode. We standardized the luminescence measurements of each compound against the distribution of DMSO controls within each assay plate. We considered a compound to be a hit if the Z-score for both screening replicates was less than or equal to −3.

To confirm the hits from the primary screen, we rescreened the compounds using the same recombinant histone peptide substrate from the primary screen, as well as a recombinant nucleosome substrate. Each screen was performed with two independent replicates as described above. Again, we considered a compound to be a hit if Z-score ≤ −3 for both screening replicates. To exclude potential false positive hits that could interfere with the AlphaLISA assay, we also screened the compounds in an AlphaLISA TruHits assay (Revvity Health Sciences cat. no. AL900D) according to the manufacturer’s directions. We considered a compound to be a hit in the AlphaLISA TruHits assay if the Z-score for both screening replicates was greater than −2.

We then prioritized 20 compounds that were hits within the secondary histone peptide, nucleosome, and AlphaLISA TruHits screens to perform dose-response assays on recombinant nucleosome substrates. We ordered the prioritized set of compounds from ChemBridge and assayed the compounds in the recombinant nucleosome assay at doses of 100 uM, 10 uM, 1 uM and 0.1 uM. The compounds were administered using a HP D300e digital dispenser and the assay was carried out as described above with two independent replicates per compound per dose. IC_50_ values were calculated based on the relative luminescence signal compared to DMSO controls.

#### Molecular docking simulations and AI-guided protein-ligand binding predictions

We utilized AI-Bind^68^ to identify putative active binding sites of small molecule ligands on KDM4 demethylases. In brief, we perturbed each amino acid trigram in the amino acid sequence of KDM4C (UniProt ID: Q9H3R0) using previously published scripts (https://zenodo.org/records/7730755) and observed changes in the AI-Bind prediction for each small molecule ligand. The SMILES and InChIKeys of small molecule ligands were derived from PubChem and used as input for AI-Bind. To validate predicted binding sites, we performed molecular docking simulations. First, the 3D ligand structures in SDF format were retrieved from PubChem. The 3D structure of KDM4C was predicted using AlphaFold3^98^ and saved in pdb format using PyMOL (v3.1.3). Molecular docking simulations were performed with AutoDock Vina implemented in SwissDock^99^ using a grid for docking that encompassed the protein domain nominated from AI-Bind. We considered the protein molecules to be rigid, whereas the ligand molecules are flexible. The docking results were then visualized in PyMOL.

